# Subfunctionalization and constrained size of the immunoglobulin *loci* in *Ambystoma mexicanum*

**DOI:** 10.1101/2022.08.03.502689

**Authors:** J Martinez-Barnetche, EE Godoy-Lozano, S Saint Remy-Hernández, DL Pacheco-Olvera, J Téllez-Sosa, H Valdovinos-Torres, R Pastelin-Palacios, H Mena-González, L Zambrano-Gonzalez, C López-Macías

## Abstract

**Background:** The axolotl, *Ambystoma mexicanum* is a unique biological model for complete tissue regeneration. Is a neotenic endangered species and is highly susceptible to environmental stress, including infectious disease. In contrast to other amphibians, the axolotl is particularly vulnerable to certain viral infections. Like other salamanders, the axolotl genome is one of the largest (32 Gb) and the impact of genome size on Ig *loci* architecture is unknown. To better understand the immune response in axolotl, we aimed to characterize the immunoglobulin *loci* of *A. mexicanum* and compare it with other model tetrapods.

**Methods:** The most recently published genome sequence of *A. mexicanum* (V6) was used for alignment-based annotation and manual curation using previously described axolotl Ig sequences or reference sequences from other tetrapods. Gene models were further curated using *A. mexicanum* spleen RNA-seq data. Human reference genomes, *Xenopus tropicalis*, and *Danio rerio* (zebrafish) were used for comparison.

**Results:** Canonical *A. mexicanum* Heavy chain (IGH), lambda (IGL), sigma (IGS) and Surrogate light chain (SLC) *loci* were identified. No kappa *locus* was found. More than half of the IGHV genes and the IGHF gene are pseudogenes, there are no clan I IGHV genes and CDRH3 diversity is restricted. Although the IGH *locus* size is proportional to genome size, we found local size restriction in the IGHM gene and in the V gene intergenic distances. In addition, there were V genes with abnormally large V-intron sizes, which correlated with loss of gene functionality.

**Conclusion:** The *A. mexicanum* immunoglobulin *loci* share the same general genome architecture as most studied tetrapods. Consistent with its large genome, Ig *loci* are larger; however, local size restrictions indicate evolutionary constraints likely to be imposed by high transcriptional demand of certain Ig genes, as well as the V(D)J recombination over very long genomic distance ranges. The *A. mexicanum* has undergone an extensive process of pseudogenization which partially explains a reduced potential repertoire diversity that may contribute to its impaired antibody response.

## Introduction

Adaptive immunity is a vital feature of the vertebrate immune system. It relies on generating a vast antigen receptor repertoire clonally distributed on the surface of B and T cells. Antigen receptor repertoire is generated independently of antigen by somatic recombining V D and J segments in primary lymphoid organs. In the periphery, antigen clonally selects B and T cells bearing high-affinity antigen receptors for the selecting antigen. In mammals, further antigen-dependent diversification is achieved by somatic hypermutation (SHM) and class switch recombination (CSR), which occur in germinal centers within the secondary lymphoid organs.

Despite the commonalities in the generation of vertebrate adaptive receptor repertoire diversity, lineage-specific evolutionary forces have shaped them differently, either by conserving the prototypic structure/recognition mode or by generating alternative and diversified structures with novel recognition and functional capabilities, so there is significant plasticity and lineage-specific rapid evolution of specific adaptive immune receptor *loci*, leading to differences in the kind of antibody classes and number variation of functional V, D and J segments involved (Das, Nozawa, et al. 2008; Das et al. 2012; Flajnik 2018).

*Ambystoma mexicanum* is a unique caudate amphibian endemic of central Mexico and is currently an endangered species. It is a neotenic organism capable of complete limb and nervous system regeneration (Voss, Woodcock, and Zambrano 2015). Like many amphibians, increased susceptibility to certain infectious diseases may play a role in axolotl population decrease (Frías-Alvarez et al. 2008; Wake and Koo 2018). Most of our amphibian adaptive immune system knowledge derives from the anurans *Xenopus leavis* and *X. tropicalis*, metamorphosing frogs (Robert and Ohta 2009). An important difference between anuran and mammalian adaptive immune systems is the former’s lack of germinal centers. Nevertheless, CSR and SHM still occur in *Xenopus* (Flajnik 2018). *Ambystoma sp*., but not other salamanders and not anuran amphibians are particularly vulnerable to certain viral infections (Charlemagne and Tournefier 1977; Charlemagne 1979, 1981; Cotter et al. 2008; Liu et al. 2014; Fonte et al. 2015). The divergence between anurans and caudate is estimated to occur 292 my ago (Pyron 2011). To gain further knowledge to make generalizations about the amphibian adaptive immune system, a thorough characterization of non-anuran species is required.

*A. mexicanum* genome has been sequenced. It is 10 times larger than the human genome (32 Gigabases) and 17 times larger than *X. tropicalis* (Nowoshilow et al., 2018). Further efforts involving SNP’s segregant mapping (Smith et al., 2019) and Hi-C allowed a 14 chromosome-level assembly (Schloissnig et al., 2021). According to the “accordion model”, genome size depends on the accumulation of transposable elements counteracted by DNA loss (Kapusta, Suh, and Feschotte 2017). The genome analysis shows that LTR retrotransposons have contributed to genome size increase in the axolotl. Moreover, lower rates of small DNA deletion compared to other tetrapods may contribute to the large genome in salamanders (Sun, López Arriaza, and Mueller 2012). V(D)J recombination is a complex mechanism that depends on non-coding transcription and deep chromatin reorganization over long DNA intervals within Ig and TCR *loci* to proceed successfully while avoiding genomic instability. If genome size affects receptor *loci* architecture and V(D)J recombination is still unknown.

To better understand the immune response in non-anuran ectotherms and explore the impact of increased genome size in adaptive antigen receptor *loci*, we report a comprehensive structural map, annotation, and functional characterization of the heavy and light chains immunoglobulin *loci* in axolotl. Moreover, we provide evidence that the IGHM gene and the IGHV and IGLV cluster size have not grown proportionally to the genome size.

## Results

### *A. mexicanum* Heavy Chain *locus* (IGH)

We identified the IGH *locus* at 36.4 - 48.5 Mbp of chromosome 13q. The whole *locus* spans 12.1 Mbp and, as in *X. tropicalis*, is flanked by the *SCL7A7* and *OXAL1* genes in the centromeric flank and the *ABDH4* and *DAD1* genes in the telomeric flank (Figure 1 A-B). We confirmed the observations suggesting that in *X. tropicalis*, the IGH *locus* is genetically linked to the TCRα/TCRδ *locus* (Parra et al. 2010). Indeed, *X. tropicalis* IGH was mapped to chromosome 1 (128.3 - 128.9 Mbp) and is only separated from the TCRα/TCRδ *locus* (128.95 - 129.5 Mbp) by the *ABDH4* and *DAD1* genes (Additional file 1: Figure S1). A detailed annotation file is provided in Additional file 2: Table S1. In contrast, the *A. mexicanum* IGH *locus* is delinked from the TCRα/TCRδ *locus*, which is found across the centromere in chr13p (3.5 - 6.5 Mbp) (Manuscript in preparation).

**Figure 1:**
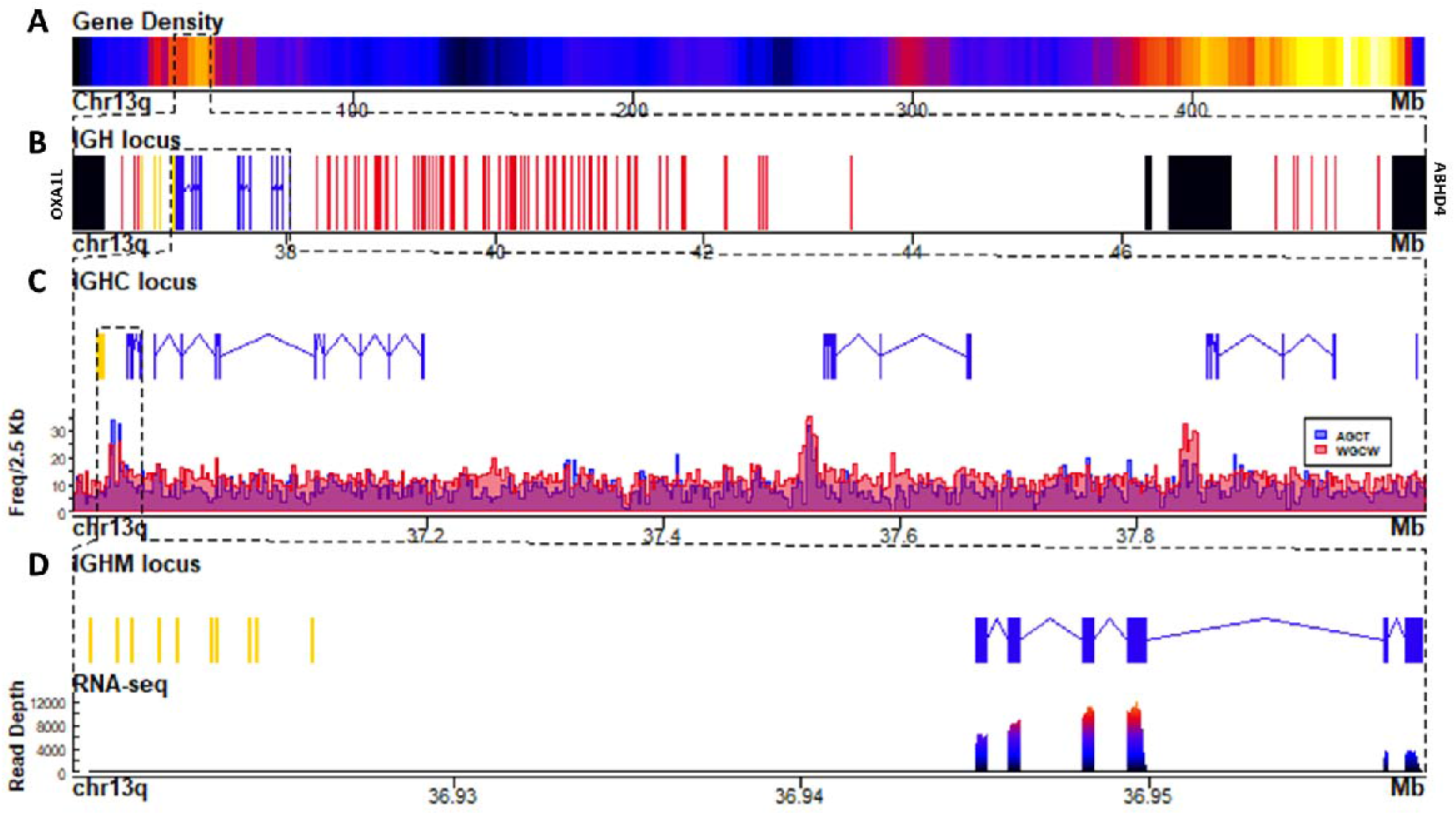
The Heavy chain *locus* is located in the centromeric portion of chr13q (483 Mbp). **A)** Gene density plot of chr13q where the IGH *locus* is encoded (box). Dark blue colors indicate low gene density. **B)** Zoom of the whole IGH *locus* (35.967 – 48.91 Mbp). Non-Ig genes (black) in proximal flank *OXA1L* gene and distal flank *ABHD4*, IGHC genes (blue), IGHV genes (red), and IGHD and IGHJ (yellow). **C)** Zoomed view (36.9 – 38.09 Mbp) of the IGHJ-IGHC gene cluster. **D)** Mapping of Switch regions using AID hotspots density per 2.5 Kb along the IGHJ-IGHC cluster. **E)** Close-up of the IGHJ cluster and the IGHM gene. **F)** Spleen RNA-seq coverage histogram of the IGHJ-IGHM region.

The usual organization of Ig and TCR *loci* is a cluster of V genes (upstream), followed by the (D) J, and C genes (downstream), all in the same transcriptional orientation. The *A. mexicanum* IGH *locus* displays an unusual organization: it starts with a 3 IGHV segment subcluster, followed by the IGHD, IGHJ, and the IGHC clusters in the same orientation (from centromere to telomere). However, downstream of the IGHC cluster is a large cluster of 85 IGHV genes delimited by the *ABDH4* and *DAD1* genes (Figure 1B). Moreover, the orientation of IGHV segments of both clusters is irregularly intercalated, with three large stretches of 19, 12, and 10 IGHV genes in the same direction, intercalated with small stretches of IGHV in the opposite transcriptional orientation (Additional file 1: Figure S2). These anomalies are likely derived from assembly errors. We propose an IGH *locus* model based on synteny with *X. tropicalis*, in which the first IGHV cluster, the IGHD, IGHJ, and IGHC conform to a single DNA block in inverted orientation so that it follows the canonical V(D)JC order in the same transcriptional orientation (Additional file 1: Figure S1).

The *A. mexicanum* IGHC cluster is 1.1 Mbp long (Chr13q: 36.94 - 38.03 Mbp), and its structure is very similar to *X. tropicalis* (Zhao et al. 2006): IGHM (4 C_H_ exons), IGHD (8 C_H_ exons), IGHX (4 C_H_ exons) and IGHY (4 C_H_ exons). All C genes encode for a transmembrane (TM) and intracellular exon (IC). The 10 exons encoding for IGHD span over 0.228 Mbp (Figure 1C).

In *X. tropicalis*, an additional C gene, IGHF, is downstream of the IGHY gene. Is composed of two C_H_ exons separated by a hinge exon, followed by TM exons (Zhao et al. 2006). We identified a region coding for a partial reading frame homologous to a single C-type Ig domain (PF07654.14), but no evidence of additional C_H_ exons or transcriptional activity within this interval, indicating that in *A. mexicanum*, IGHF is a pseudogene (Figure 1C; Additional file 1: Figure S1).

Class switch recombination is initiated by cytidine deamination mediated by the Activation Induced Cytidine Deaminase (*AICDA*), which preferentially uses the 5’-AGCT-3’ motif as a deamination hotspot (Xu et al. 2012). Based on the density of occurrence of such motif (Figure 1C), we identified two highly enriched regions (z-score > 4) (Additional file 1: Figure S3 A-B). As expected, the first corresponds to the Sμ region and is located within the 36.93 - 36.942 Mbp interval between the IGHJ cluster and the first IGHM exon. The second region corresponds to the Sχ region and was located within the 37.517 - 37.532 Mbp interval, upstream of the first IGHX exon (Figure 1C; Additional file 1: Figure S3C).

No 5’-AGCT-3’ motif enrichment was found upstream of the IGHY or IGHF pseudogene (Figure 1C; Additional file 1: Figure S3D). Interestingly, a conspicuous enrichment of the 5’-RGYW-3’ in the direct strand and the 5’-WGCW-3’ palindrome was located in the 37.837 - 37.852 Mbp interval upstream of the IGHY first exon, which corresponds to the Sυ region (Additional file 1: Figure S3D). Further characterization of RGYW motifs revealed that the ACTG motif is also abundant in Sμ but depleted in Sυ. The AGTA motif was abundant in Sυ (predominantly in the reverse strand) and depleted in Sμ and Sχ (Additional file 1: Figure S4 A-B). Together, these results indicate that the composition of AID targets may vary within S regions. Except for the absence of the Sφ region, the occurrence of switch regions follows a similar pattern as in *X. tropicalis* (Zhao et al. 2006).

The IGHJ cluster is composed of nine functional IGHJ segments and one pseudogene. Functional IGHJ segments encode for the canonical WGXG motif and have a 23-bp spacer and highly conserved heptamer and nonamer in their J-RSS. The pseudogene (IGHJ10) is the most proximal to IGHM and lacks a conserved heptamer and nonamer in the RSS (Additional file 1: Figure S5).

Within the 36.6 - 36.8 Mbp interval, upstream of the IGHJ *locus*, we found the IGHD *locus* composed of only four IGHD genes, flanked in both directions by 12 bp spaced RSS’s. The length of three IGHD genes was 11 bps, and one was 13 bp. All IGHD genes were G+C rich, lacked stop codons in the 6 reading frames, and showed partial identity to previously described IGHD core sequences (Golub, Fellah, and Charlemagne 1997) (Additional file 1: Figure S6). To gain insight into CDRH3 junctional diversity, we sequenced the IgM cDNA repertoire by targeted sequencing of 5’RACE amplicons derived from the spleen of three adult individuals (Figure 2A). Some translated frames of IGHD_02, 03, and 04 were found in CDRH3 junctions previously described and our own Rep-seq data, confirming their authenticity as IGHD genes. Accordingly, with IGHD gene lengths, CDRH3 length in axolotl was shorter than human (Figure 2B), and there was frequent clonotype sharing within individuals indicating a less diverse repertoire (Figure 2C).

**Figure 2:**
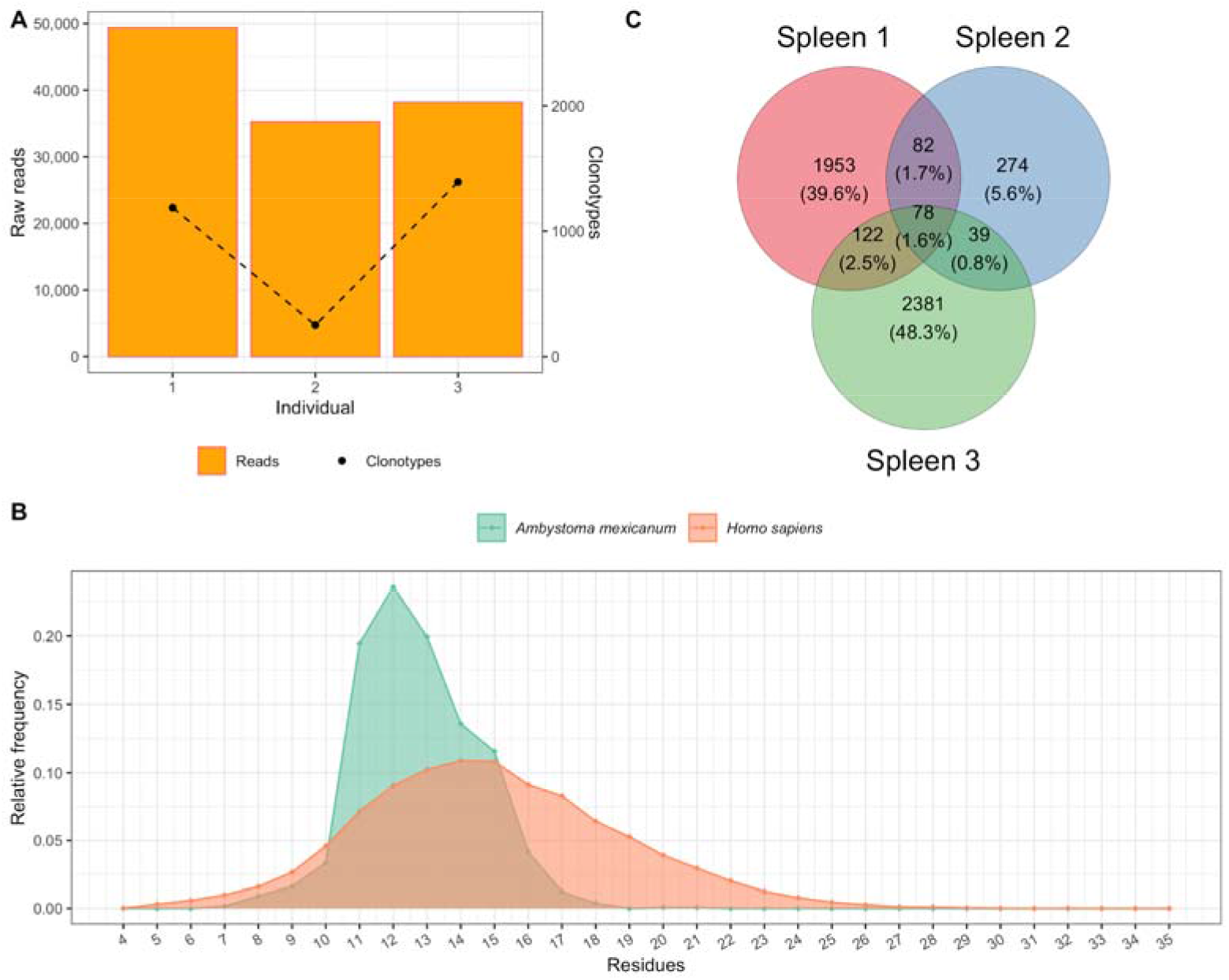
Rep-seq analysis of *A. mexicanum* heavy chain variable region from spleen mRNA. **A)** Number of paired sequencing reads and VH clonotypes identified from each individual. **B)** Pooled CDRH3 length distribution in *A. mexicanum* IgM compartment from the spleen of three individuals and comparison with healthy human IgM CDRH3 length distribution. **C)** Number of shared spleen IgM HC clonotypes between three individuals.

We found 99 IGHV genes, of which 88 map in chr13q, and the remaining are located in unmapped scaffolds (Table 1). Based on the analysis of the functionality of the coding sequence, the presence of RSS’s and transcription by RNA-seq and Rep-seq, we classified 47 IGHV genes as functional, 22 as ORF’s and 30 as pseudogenes (Additional file 2: Table S2). Three IGHV segments (IGHV_083, IGHV_084, and IGHV_085) are non-functional pseudogenes located outside the IGH *locus*, specifically within the class II MHC *locus* in the telomeric region of chr13q (464.8 - 465 Mbp) (Additional file 2: Table S1) (Schloissnig et al. 2021). As expected, most IGHV segments had a recognizable 23 - bp spacer in their 3’ RSS. However, three apparently functional IGHV segments mapped to chr13q (IGHV_034, 035, and 055) had a 12 - bp spacer between their apparently functional RSS, so we labeled them as ORF’s for violating the 12/23 rule (Additional file 2: Table S1).

**Table 1.** Summary of variable genes per *loci*.

|  |  | Functional | (%) | ORF | (%) | Pseudogene | (%) | Total |
| --- | --- | --- | --- | --- | --- | --- | --- | --- |
| **IGH**<br>*locus* | **Chr13q**<br>**non-mapped** | 43 | (43) | 20 | (20) | 25 | (25) | 88 |
|  |  | 4 | (4) | 1 | (1) | 6 | (6) | 11 |
|  |  | 47 | (47) | 21 | (21) | 21 | (31) | 99 |
| **λ**<br>*locus* | **Chr10p**<br>**non-mapped** | 44 | (62) | 5 | (7) | 21 | (30) | 70 |
|  |  | 1 | (1) | 0 | (0) | 0 | (0) | 1 |
|  |  | 45 | (63) | 5 | (7) | 21 | (30) | 71 |
| **σ**<br>*locus* | **Chr 1p**<br>**non-mapped** | 5 | (83) | 1 | (17) | 0 | (0) | 6 |
|  |  | 0 | (0) | 0 | (0) | 0 | (0) | 0 |
|  |  | 5 | (83) | 1 | (17) | 0 | (0) | 6 |
| **κ**<br>*locus* |  | Not found. Syntenic non-Ig genes found in chr6q: 1.4-1.5 Gb |  |  |  |  |  |  |

IGHV segments in tetrapods are classified in three major phylogenetic clans based on sequence conservation of the framework regions (Schroeder et al., 1990; Das et al., 2008b). Phylogenetic analysis of axolotl functional IGHV using human and mouse IGHV sequences as reference revealed 24 segments belonging to clan II, 19 segments belonging to clan III, and the absence of clan I segments (Additional file 1: Figure S7). Four IGHV genes (IGHV_070, 071, 077 and 082) could not be assigned to a particular clan (Additional file 2: Table S1). IGHV ORF’s also were assigned to clan II (n = 11) and clan III (n = 4), but none to clan I, and seven could not be assigned to any clan. Of the eleven IGHV families described by (Golub and Charlemagne 1998), we found functional representatives for all but family VH6 and VH9 (Additional file 2: Table S1). Four genes, IGHV_005, IGHV_039, IGHV_063, and IGHV_065, which belong to the VH8 family, contain a triple Cys in the CDRH1, which appears to be a unique feature of urodeles (Fonte et al. 2015). Moreover, seven IGHV genes (IGHV_051, 056, 057, 058, 061, 062, and 082) could not be assigned to any family (Additional file 1: Figure S8).

### *A*.*mexicanum* light chain *loci*

We found three Ig light chain *loci*: A large 9 Mbp *locus* in chr10p:116.7 - 125.8 Mbp (Figure 3), a 0.1 Mbp *locus* at chr1p:609.1 - 609.2 Mbp (Figure 4 A-E), and a third 0.2 Mbp *locus* at chr1p: 68.0 - 68.2 Mbp (Figure 4 A-B).

**Figure 3:**
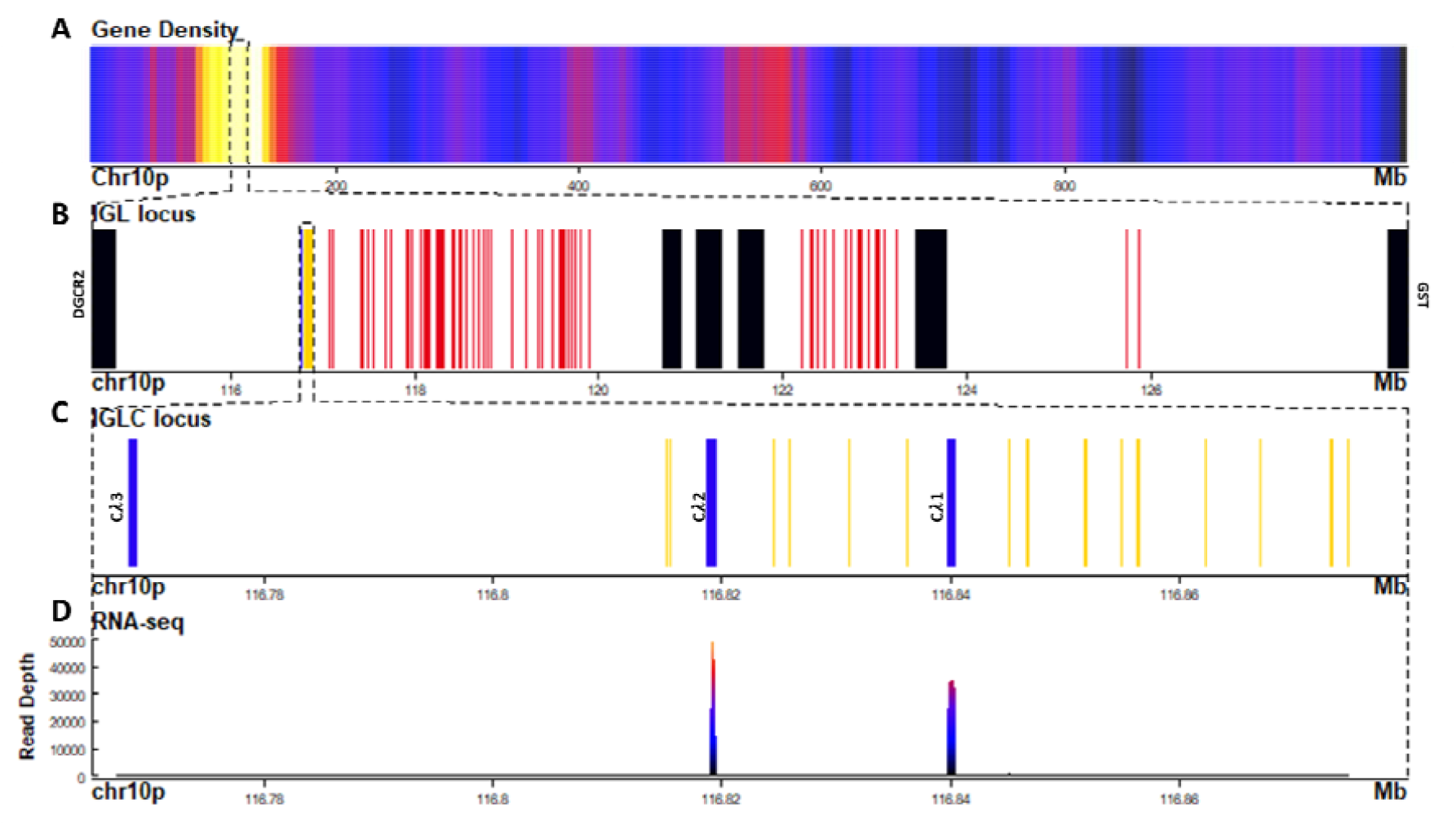
The lambda light chain *locus* is located in the centromeric portion of chr10p (116.7 - 125.8 Mbp). **A)** Gene density plot of chr10p where the IGL *locus* is encoded (box). Dark blue colors indicate low gene density. **B)** Zoom of the whole IGL *locus* (114.5-128.8 Mbp). Non-Ig genes (black) in proximal flank DGCR2 gene, and distal flank GST. Zoomed view of IGLC genes (blue), IGLV genes (red), and IGLJ (yellow). **C)** Zoomed view (116.76-116.88 Mbp) of the IGHC gene cluster. **D)** Spleen RNA-seq coverage histogram. Note that there is no expression in Cλ3

**Figure 4:**
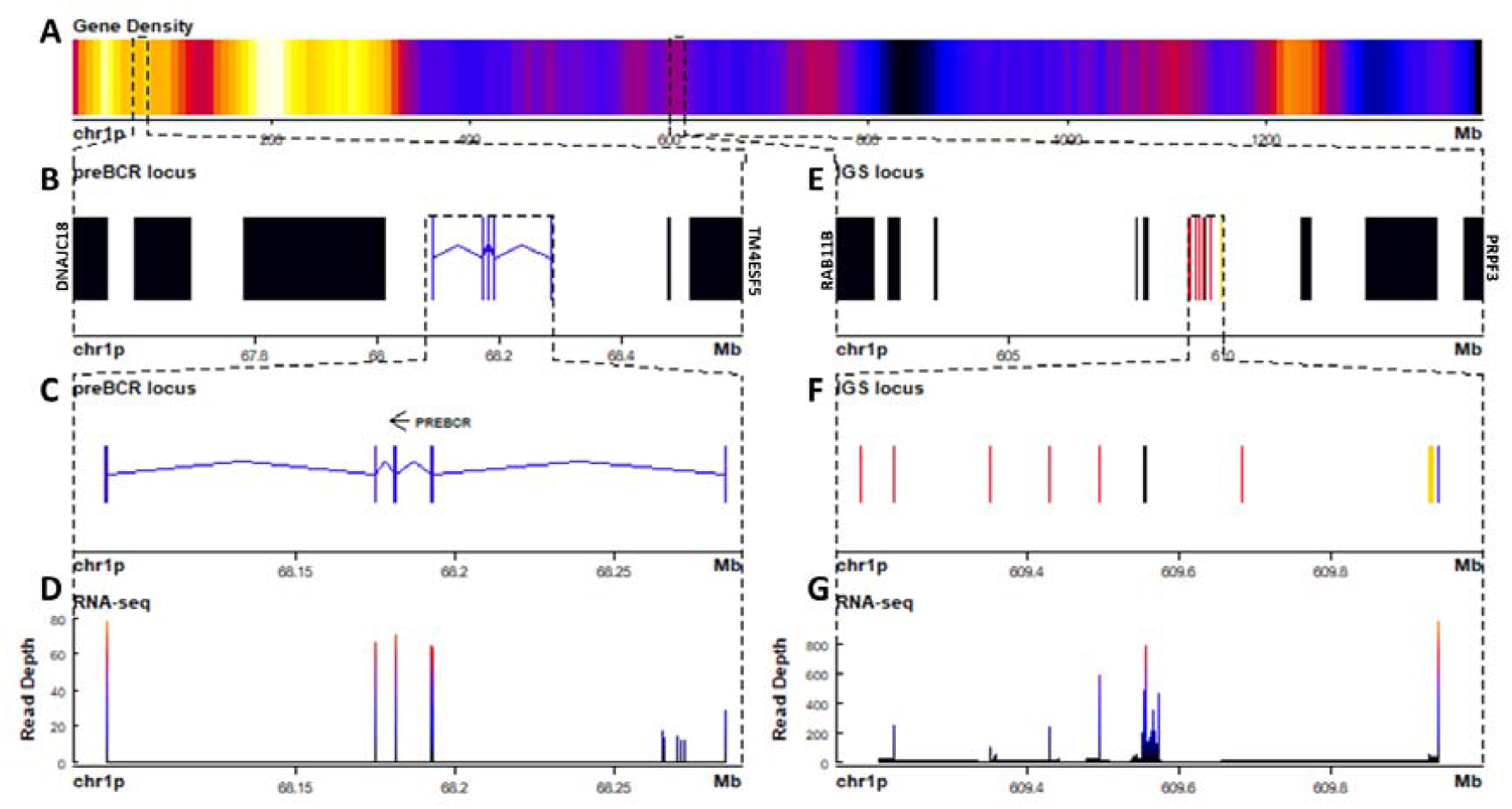
The sigma light chain *locus* is located in the centromeric portion of chr1p (609.1 - 609.2 Mbp). **A)** Gene density plot of chr1p where the IGS and VpreB *loci* are encoded (box). Dark blue colors indicate low gene density. **B)** Zoom of the whole VpreB *locus* (67.5-68.6 Mbp). Non-Ig genes (black) in proximal flank *DNAJC18* and distal flank *TM4ESF5*. Zoomed view of IGLC genes (blue). **C)** Zoomed view (68.08-68.29 Mbp) of the IGSV and IGSC gene cluster. **D)** Spleen RNA-seq coverage histogram. **E)** Zoom of the whole IGS *locus* (601-616 Mbp). Non-Ig genes (black) in proximal flank *RAB11B* gene and distal flank PRPF3. **F)** Zoomed view of IGSC genes (blue), IGSV genes (red) and IGSJ (yellow) (609.15-610 Mbp) **G)** Spleen RNA-seq coverage histogram.

The larger *locus* in chr10p corresponds to the lambda (IGL) and is similarly structured as in *X. tropicalis*, also referred to as the type III light chain *locus* (Haire et al. 1996; Qin et al. 2008). It is composed of a tandem of three Cλ genes and their corresponding IGLJ gene clusters, followed by a single IGLV cluster (Figure 3 A-B). The three Cλ genes encode for the characteristic Lys23 and Lys60 unique to Cλ and cluster together with human and *X. tropicalis* Cλ protein sequences (Figure 3C; Additional file 1: Figure S9). Only Cλ1 and Cλ2 were transcribed in spleen RNA (Figure 3D).

IGLJ cluster 1 (upstream of Cλ1) is the largest cluster with 9 IGLJ gene segments (IGLJ1.1-1.9), all functional but IGLJ1.6, which lacks a conserved heptamer at the RSS. The second IGLJ cluster comprises four functional IGLJ segments (IGLJ2.1-4). The third cluster comprises only two functional IGLJ segments (IGLJ3.1-2). Eleven out of 15 IGLJ segments encode for the canonical motif FGXG characteristic of IGLJ and IGKJ. In the remaining four, the Phe is replaced by Ile (IGLJ2.1 and 3.1) or Leu (IGLJ1.1 and 2.4) (Additional file 1: Figure S10). In contrast with IGKJ or IGSJ gene segments, all IGLJ are flanked by 12 bp-spaced RSS’s (Additional file 1: Figure S10 and S11).

The cladistic marker analysis (Das, et al. 2008) confirmed the identity of the IGLV segments, which includes the sequence gap between position 7 (FWR1), presence of gaps at positions 41a, 46a, and 46b (FWR3), presence of Ala53 instead of S/T (FRW3) and the DEAD motif in positions 64-67. Additionally, IGLV gene segments were flanked by 23 bp - spacer RSS (Additional file 1: Figure S12). The IGLV cluster comprises 71 IGLV genes, of which 45 appear functional (Table 1; Additional file 2: Table S3). Phylogenetic analysis revealed that 41 functional segments belong to the four previously described IGLV families (André et al. 2000), but IGLV_040, IGLV_069, and IGLV_037 are likely to belong to a novel family (Additional file 1: Figure S13).

As in the IGH *locus*, many IGLV segments are intercalated in opposing directions. In *X. tropicalis*, the IGL *locus* is flanked by the *DGCR2* gene on the centromeric flank and the *OTUB* genes in the telomeric flank. While in axolotl *DGCR2* also flanks IGL on the centromeric end, *OTUB, MMP11*, and *SMARCB11* split the IGLV cluster in two, but overall synteny (*SLC2A11,GST*, and the *CABIN1* genes) is maintained in the telomeric flank (Figure 3; Additional file 1: Figure S14).

The second light chain *locus* is composed of only 6 V genes (5 functional), flanked by a 12 bp-spaced RSS, 5 J genes flanked by a 23 bp-spaced RSS, and a single C gene composed of 2 exons (Figure 4 E-F; Table 1; Additional file 2: Table S1, Table S4). Is similarly structured to the *X. tropicalis* IGS *locus* (Haire et al. 1996; Qin et al. 2008), including synteny in one flank (*ANGPTL4, RAB11B*, and *MARCHF2* genes) (Additional file 1: Figure S15). Cladistic analysis is consistent with the identity of this *locus* as the σ *locus*, such as the presence of Thr7 (FWR1), absence of gap in positions 41a and 46b (FWR3), the presence of Tyr53, and absence of DEAD motif (64-67) (FWR3) in V genes, as well as the presence of FSXS motif instead of the FGXG (di-Glycine bulge) motif of IGKJ or IGLJ.

The third light chain *locus* contains a single gene composed of five exons encoding for a 307 residue polypeptide. The first exon encodes for a signal peptide, exon 2 encodes for an Immunoglobulin V-set domain (PF07686), exon 3 encodes for an Immunoglobulin Cλ-set domain (PF07654), and exons 4 and 5 encode for the C-terminal region and the 3’ UTR, with no homology identified by BLASTP to the nr NCBI database. Based on the predicted protein and gene structure, we suggest that this gene encodes for the surrogate light chain (SLC) of the preB cell receptor.

In *X. tropicalis*, a substantial portion of the light chain repertoire is composed of κ light chains, also referred to as □ (rho) *locus* (Qin et al. 2008). It is encoded in the telomeric tip of chr1 (2-2.8 Mbp). We did not find evidence of the κ *locus* in *A. mexicanum*, although non-Ig the orthologs in the centromeric flank (*SUCLG1)* and telomeric flank (*ADRA1D* and *RNF24*) of the *X. tropicalis* κ *locus* were found in *A. mexicanum* chr6: 1444 – 1503 Mbp (Additional file 1: Figure S16).

### Implications of genome size in the architecture of IGH *locus* in tetrapods

The *A. mexicanum* genome is 18.8, 10.3, and 21 times larger than the *X. tropicalis* (Hellsten et al. 2010), human (Piovesan et al. 2019) and zebrafish genome (Driever et al. 1994), respectively. We asked if the IGH *locus*, the IGHC cluster, and individual IGHC gene lengths were proportional to their genome size difference. The axolotl IGH *locus* was 22, 9.4 and 70 times larger than the *Xenopus*, human, and zebrafish *locus*, respectively, which in the case of *Xenopus* and human is roughly proportional to the corresponding genome size (Additional file 2: Table S5). The IGHC in *A. mexicanum* was 10, 4, and 22 times larger than in *X. tropicalis*, human, and zebrafish, respectively. Interestingly, while IGHD, IGHX, and IGHY were 8 - 12 times larger in *A. mexicanum* than in *X. tropicalis*, IGHM was only 1.4 times larger (Additional file 1: Figure S17; Additional file 2: Table S5). Similarly, *A. mexicanum* IGHM was 2.9 times larger than the human and zebrafish ortholog, whereas the IGHD was Table S625 and 18 times larger, respectively. These results suggest that some regions of the IGH *locus*, and particularly the IGHM, but not IGHD gene intron length, is evolutionarily constrained.

### V-intron length

The first exon of each V gene encodes for a signal peptide that allows the BCR or TCR to follow the secretory pathway. We reasoned that genome enlargement by genome drift would be accompanied by increasing V-intron lengths unless intron size is functionally constrained. By measuring the distribution of V-intron length, we observed that the V introns of functional *A. mexicanum* IGHV genes were shorter than in IGHV pseudogenes (P = 0.037; Figure 5A). No differences in V intron length were noted between functional and non functional IGLV genes (P = 0.84; Figure 5B) or IGHV introns in *D. rerio* (P = 0.45; Figure 5C), *H. sapiens* (P = 0.44; Figure 5D), and *X. tropicalis* (P = 0.13; Figure 5E). However, comparison of *A. mexicanum* V-intron length IGHV across species revealed no differences in functional IGHV genes (P > 0.05; Figure 5F), whereas non-functional IGHV of *A. mexicanum* were larger than human (P = 0.009), zebrafish (P = 0.003) and frog (P = 0.002) (Figure 5G; Additional file 2: Table S6).

**Figure 5:**
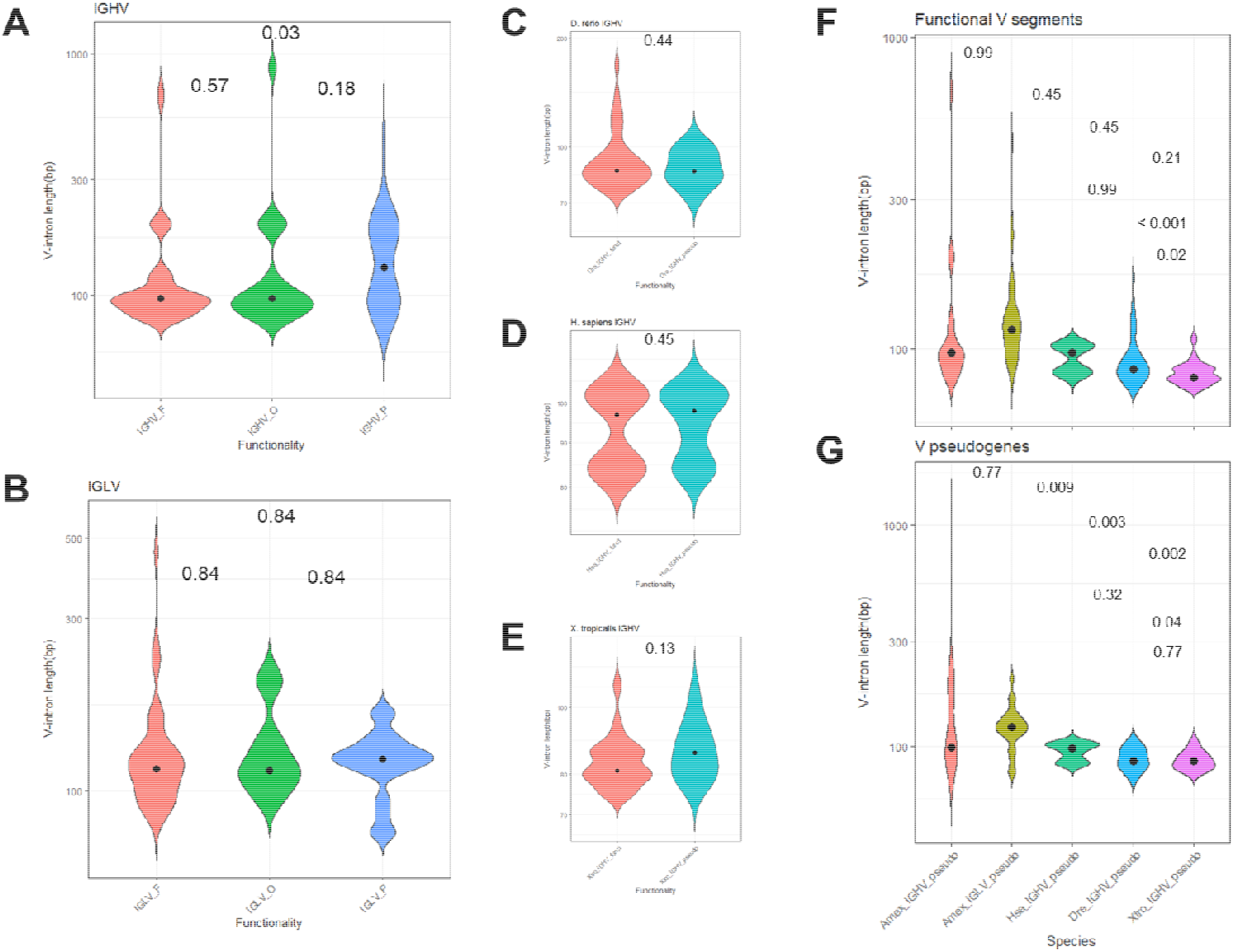
Distribution of V intron length in tetrapods. The violin area is scaled for comparability. The median is shown as a black dot. **A)** the heavy chain and **B)** lambda light chain *locus* in *A. mexicanum* according to their functionality class (Functional, ORF, pseudogene). In *A. mexicanum*, Intron length in functional IGHV genes was shorter than in IGHV pseudogenes. No differences were observed in the lambda *locus* and in **C)** *D. rerio*, **D)** *H. sapiens*, and **E)** *X. tropicalis*. V Introns in functional IGHV *A. mexicanum* were not larger than functional *A. mexicanum* IGLV genes, or **F)** functional IGHV of other tetrapods, however, V introns in **G)** non-functional *A. mexicanum* IGHV genes were larger than other tetrapods. Statistical analysis for multiple comparisons **(A, B, F, and C)** was performed with the Robust ANOVA One-Way Trimmed Means Comparisons, with a trimming level of 5% (tr = 0.05), followed by the *post hoc* Lincon test (Mair and Wilcox, 2020). “Robust Statistical Methods in R Using the WRS2 Package.” Behavior Research Methods, 52, 464–488) (Mair and Wilcox 2020). For two sample comparisons (C-F), Yuen’s robust Tests for Two Independent Groups were performed with a trimming level of 5 %.

To further explore a relationship between V-intron length and functionality in *A. mexicanum*, we tested if the observed number of long introns (> 150 bp) were more frequent in non-functional IGHV and IGLV segments than expected by chance. A Fisher’s exact test revealed that the odds of an IGHV gene with a long V-intron being non-functional are 3.3 higher than its short V-intron counterpart (P = 0.018, CI95: 1.1, 10.7) (Additional file 1: Figure S18A). In contrast, for the IGLV *locus*, there were no differences in the observed and expected frequencies of short and long V-introns according to functionality (P = 0.73; OR = 0.63, CI95: 0.09, 3.0). Overall, these results indicate that in *A. mexicanum*, an increase in V-intron length is evolutionarily constrained in IGHV but not IGLV (Additional file 1: Figure S18B).

### IGHV and IGLV intergenic lengths are shorter

We reasoned that an increase in *A. mexicanum* genome size would not impact the Ig *loci* architecture unless there are functional constraints for Ig *loci* enlargement. The size difference of the IGHV gene cluster in *A. mexicanum* (17, 7, and 70 times larger than *X. tropicalis*, human, and zebrafish, respectively) is roughly proportional to its genome size (Additional file 1: Figure S18; Additional file 2: Table S6). We further compared functional IGHV and IGLV intergenic length distribution in *mexicanum, H. sapiens, D. rerio*, and *X. tropicalis*. As a reference for comparison, we used the whole genome coding genes and cytochrome *p450* family (a non-Ig gene family commonly encoded in gene clusters) intergenic distance. As expected, *A. mexicanum* whole-genome (Figure 6A), IGHV and IGLV (Figure 6B), and *p450* intergenic lengths (Figure 6C) were larger than their corresponding distributions in all tested species. In *A. mexicanum*, IGLV intergenic lengths were larger than IGHV (Figure 6B). Whole-genome intergenic lengths were larger than IGHV (P = 2.4e-22) and *p450* (P = 3.7e-04), and IGHV were smaller than *p450* (P = 1.0e-03) (Figure 6D). Although *p450* intergenic lengths in *H. sapiens* and *D. rerio* were no different than whole-genome (Figure 6 F-G), *A. mexicanum* and *X. tropicalis* were smaller (Figure 6 D-E). Nevertheless, we observed that in all species, IGHV intergenic lengths were consistently smaller than the whole-genome and *p450* (Figure 6 D-G), indicating that the IGHV cluster size is constrained by natural selection.

**Figure 6:**
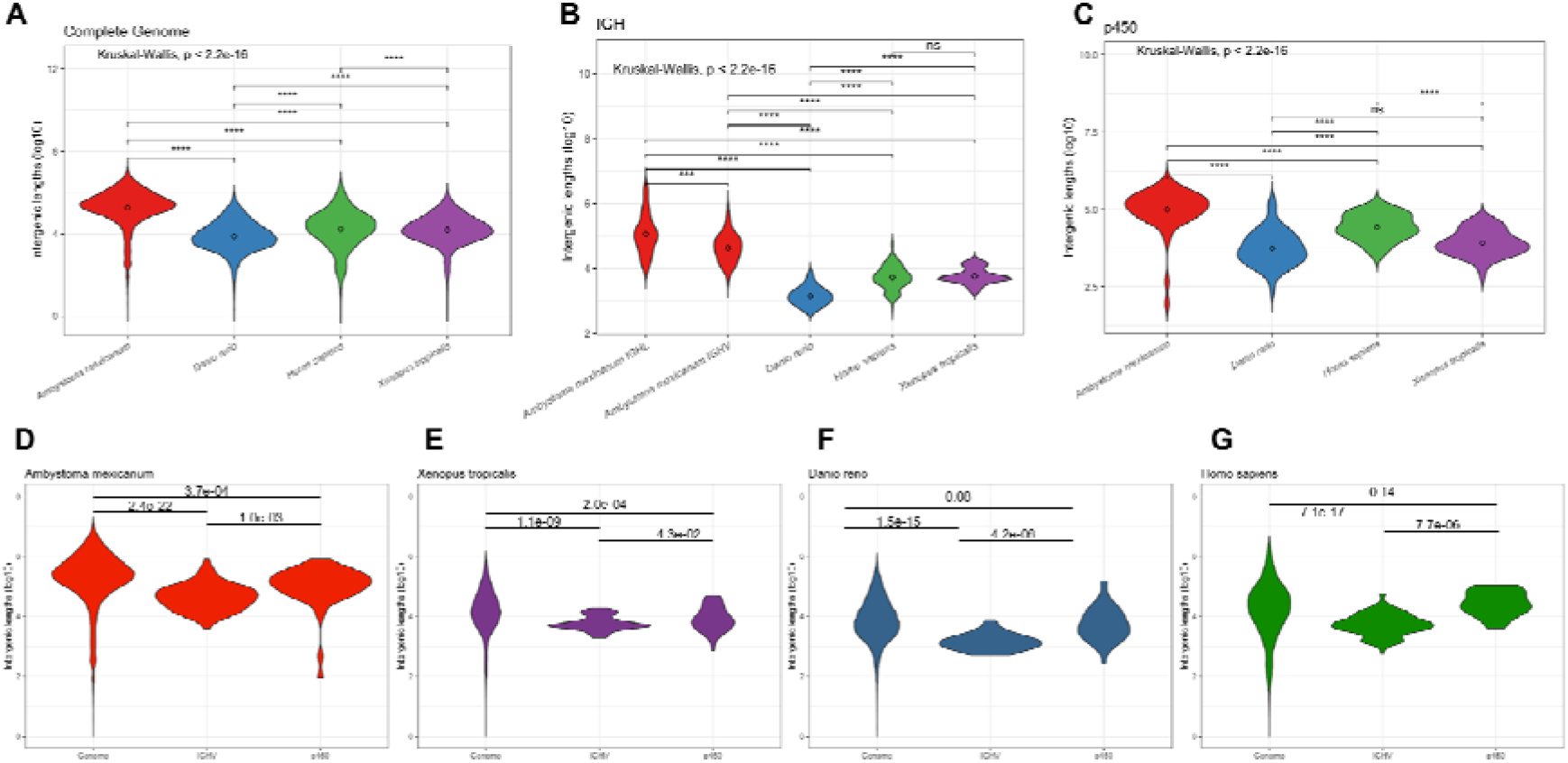
Intergenic distances by species. Violin plots depicting overall coding to coding gene intergenic distance distribution across the whole genome (genome), between IGHV segments, and between cytochrome *p450* family gene clusters in *A. mexicanum* (red), *X. tropicalis* (purple), *D. rerio* (blue), and *H. sapiens* (Green). Kruskal-Wallis rank sum test followed by Dunn’s multiple comparison test with Benjamini – Hochberg p adjustment.

## Discussion

The availability of a high-quality genome draft of *A. mexicanum* assembled at the chromosome level enabled us to evaluate Ig *loci* organization and its relation to previous functional analysis of the axolotl humoral immune response (Fellah and Charlemagne 1988; Golub, Fellah, and Charlemagne 1997; Golub and Charlemagne 1998; Smith et al. 2019). Moreover, it provides the opportunity to investigate if the Ig *loci* grow in parallel with the rest of the genome or if there are functional constraints that limit Ig *loci* enlargement.

From a comparative immunogenomic perspective, we found loss of function of the IGHF gene, limited VH junctional diversity, lack of IGHV segments belonging to the tetrapod clan I, and loss of the IGK *locus*. From a genome evolution perspective, we found an unusual increase of V-intron size in a subset of IGHV and IGLV segments and that although Ig *loci* size has increased, the IGHM gene and the IGHV and IGLV intergenic distances have not grown at the same rate as the rest of the genome.

We identified an atypical configuration of the IGH *locus*, particularly regarding the position of the IGHJ and IGHC genes upstream of the IGHV cluster (Figure 1). This unusual architecture would be incompatible with B cell maturation and antibody production because V(D)J recombination involving IGHV segments downstream IGHJ *locus* would delete the IGHC cluster. In the case of the IGL *locus*, VJ recombination involving the more distal IGLV segments would delete the *OTUB, SMARCB11, MMP11*, and the *SLC2A11* gene cluster. Moreover, V gene clusters in alternating coding directions would be incompatible with chromatin extrusion during V(D)J recombination due to frequent convergence of CBE’s (Zhang et al. 2022) (see below). We propose that such inconsistencies may be the result of local scaffold orientation errors attributable to the inherent complexity and genome size, combined with the repetitive nature of the Ig/TCR *loci*, despite the enormous technological effort involving long-read sequencing (Nowoshilow et al., 2018), SNP segregant and fiscal mapping (Smith et al. 2019), and Hi-C mapping (Schloissnig et al. 2021).

Apart from such inconsistencies, the overall organization of the Ig *loci* in *A. mexicanum* is similar to the anuran *X. tropicalis*, and we confirmed studies describing the main antibody classes, IgM and IgY (Fellah, Jacques, and Charlemagne 1994), IgX (Schaerlinger and Frippiat 2008) and IgD (Yoshinaga et al. 2021). Interestingly we demonstrate that IGHF is a pseudogene. IGHF in *Xenopus* is one of the earliest examples of an antibody hinge region encoded by an exon (Zhao et al. 2006). The functional implications of IGHF loss of function axolotl are unknown but worthy of further research.

IgM repertoire sequencing of spleen mRNA allowed us to refine IGHV models but revealed a restricted CDRH3 size distribution due to the shortness of only four IGHD segments. VH clonotype analysis revealed a high proportion of shared clonotypes within 3 individuals, confirming a reduced junctional diversity (Golub and Charlemagne 1998).

Studies by Charlemange’s group anticipated that most if not all light chains in axolotl were λ chains (André et al. 2000). Extensive homology-based sequence searches in the previous genome version (V3) and the current version (V6) revealed the absence of the IGK or □ *locus* in axolotl. Search for cladistic markers, RSS configuration, and overall *locus* architecture allowed us to unambiguously define that the only light chain *loci* identified in current assemblies correspond to λ, σ, and the Surrogate light chain (SLC), a component of the preB cell receptor. Genome search for IGK in other caudate amphibians may reveal if IGK loss is restricted to axolotl.

Larger introns may be biologically costly by increasing the investment of nucleotides and time, compromising the fidelity of the resulting mRNA, and increasing the space for allelic variation and aberrant splicing (Lynch, 2007). Recent *in situ* studies in human cells show that splicing efficiency is not affected by exon number and intron lengths (Singh and Padgett 2009); (Hollander et al. 2016). Comparative analysis of intron length distribution in axolotl revealed that introns of genes involved in developmental processes are shorter than non-developmental genes, suggesting an evolutionary constraint that favors higher transcriptional rates (Nowoshilow et al. 2018). Similarly, we observed that the IGHM gene size in axolotl, humans, *Xenopus* and zebrafish is very stable despite the wide variation in genome size. As the size of the IGHM coding exons is constant, our results indicate an intron size constraint favoring IGHM transcription levels or regulation.

Intron size also varies within each gene, and constitutive exons are flanked by shorter introns than alternatively spliced exons (Gelfman et al., 2012). The first exon of each V gene encodes for the majority of the signal peptide (L-PART1) in immature H or light chain polypeptides and, as such, are constitutive exons. Except for a subset of abnormally large V-introns found in the axolotl (up to 800 bp), we found that V-intron length is usually short (80-100 bp), as in the remaining species tested. Also, it is more likely that IGHV pseudogenes have abnormally large V-introns than their functional counterparts. Moreover, only in the axolotl, V-intron length is larger in IGHV pseudogenes than in functional IGHV genes. However, this association was not found in the lambda *locus*. Furthermore, the surrogate light chain is a non-recombined light chain that associates with successfully recombined μ chains and is expressed only during proB to preB cell development at relatively low levels. Noteworthy, in contrast to human and mouse where the SLC is encoded by two genes, VPREB and λ5 (Melchers 1999), the SLC is axolotl is encoded by a single gene. The axolotl SLC gene spans over 195 Kb, and the V-intron is 93 Kb long. Assuming a similar RNApolII elongation rate to humans (Singh and Padgett 2009), an SCL RNA precursor would take 51 min to synthesize. We propose that V-intron lengths at the IGH *locus*, but not the light chain and SLC *loci*, are constrained by evolution to favor stringent transcriptional control during preB cell development.

The hallmark of adaptive immunity is the generation of a vast array of clonally distributed antigen receptors generated by V(D)J recombination. Much knowledge has been generated that explains how the RAG recombinase mediates precise double-stranded breaks in V(D)J segments over tens to hundred kilobases apart. Non-coding transcription and chromatin remodeling are crucial (Schatz and Ji 2011). More recently, it has been shown that the mouse IGH *locus* is a topologically associated domain (TAD) organized into three chromatin loops or sub-TAD’s required for adequate spatial approximation of IGHV genes to the recombination center by chromatin loop extrusion (Kenter, Watson, and Spille 2021; Zhang et al. 2022). Chromatin extrusion is mediated by the cohesin ring protein complex by extruding chromatin between convergently oriented CTCF-binding elements (CBE’s). Whole-genome TAD analysis in axolotl revealed that TAD size increase does not affect function negatively (Schloissnig et al., 2021). We asked whether the size of the IGHV gene cluster is constrained in axolotl. Despite the large size of the IGHV gene cluster (6.7 Mbp), we observed that intergenic IGHV distances are significantly smaller than all genome intergenic distances or *p450* gene clusters, indicating that larger distances impose a functional cost that it does not affect non-coding transcription (required for the initiation), could affect V(D)J recombination *per se* by favoring spurious recombination events due to cryptic RSS’s.

In conclusion, *A. mexicanum* Ig *loci* share the same structure with other tetrapods, however, it shows multiple signs of subfunctionalization such as a large proportion of V pseudogenes and an IGHF pseudogene, loss of the κ *locus*, and a very small σ *locus*. We provide evidence suggesting an increase in genome size may be causally related to this subfunctionalization, particularly in the IGH *locus*, and that Ig *loci* enlargement, particularly at the IGH *locus*, has been negatively selected, likely to counteract a negative functional effect in transcriptional regulation and/or V(D)J recombination. Systematic functional immunocompetence analysis in amphibians and other vertebrates with large genomes may provide a definitive answer.

## Methods

### *A*.*mexicanum* genome data

The latest version genome sequence of *A. mexicanum* white d/d strain AmexG_v6.0-DD, and the corresponding annotation file AmexT_v47-AmexG_v6.0-DD.gtf.gz was downloaded from https://www.axolotl-omics.org/assemblies (Schloissnig et al. 2021).

### Immunoglobulin *loci* mapping

We used reference IGHV, IGHC, IGLC cDNA sequences obtained IMGT (http://www.imgt.org/) to map the IGH and IGL *loci* using TBLASTX, Exonerate (EST2genome alignment model). Significant hits (e-value < 1.0E^-05^ for BLASTX, score > 100 for Exonerate) were exported as GFF3 files to load them in the Integrative Genomics Viewer (IGV) (Thorvaldsdottir, Robinson, and Mesirov 2013) for visualization and manual curation. Searches were further complemented using HMMER3 option *hmmsearch*, using PFAM immunoglobulin domain *hmm* models (V-set: PF07686.17 and C-set: PF07654.14).

Gene models were refined by mapping transcriptomic data from *A. mexicanum* spleen from two sources: 1) Public RNA-seq spleen data (project SRP101842, run SRR5341570), downloaded from https://www.ncbi.nlm.nih.gov/sra. 2) Rep-seq data from *A. mexicanum* spleen total RNA, obtained from the UNAM strain. For mapping RNA-seq reads from both sources, we used the STAR aligner (Dobin and Gingeras 2015), using only relevant chromosome arms as targets.

We performed a new *X. tropicalis* IGH *locus* annotation based on the corresponding ENSEMBL annotation (Xenopus_tropicalis_v9.1, GCA_000004195.3), further complemented and refined with sequence data kindly provided by Sibayshi Das (Das, Nozawa, et al. 2008), and *X. tropicalis* liver RNA-seq data (SRR579561) (Barbosa-Morais et al. 2012) following the same methodology as for *A. mexicanum*.

### Definition of V, D, and J functionality

Functionality was defined based on the IMGT criteria (Lefranc 2014). For IGHV segments to be functional (F), the presence of a predicted *in-frame* signal peptide exon/V exon open reading frame (ORF) compatible with a V-domain, the presence of Cys23, Trp41, Trp 52, and Cys104 in the V exon, followed by a *bona fide* RSS (as described above), as well as transcription based on spleen RNA-seq data or AIIR-seq. We consider as non-functional V-ORF (O), the presence of a V-domain ORF in absence of Cys22, Trp36, Trp 47, or Cys104, or the absence of RSS. Frame-shifted V-ORF, interruptions by stop codons, or lack of signal peptide exon was regarded as a V-pseudogene (P).

### Phylogenetic analysis of IGHV and IGLV segments

For the heavy chain *locus, H. sapiens* and *M. musculus* functional IGHV segments were retrieved from the IMGT (Manso et al. 2022). For *X. tropicalis* functional IGHV segments were retrieved from our own annotation. Multiple sequence alignment with the corresponding *A. mexicanum* IGHV segments was performed with MUSCLE (Edgar 2004). Principal Component Analysis (PCA) of the aligned sequences was done with JalView 2.10.5 (Waterhouse et al. 2009) using a PAM250 scoring matrix.

For the comparative phylogenetic analysis of the IGL *locus*, the approach was identical as for IGHV, but using sequence data of supplementary file SD1.xls from twelve tetrapods (Das, Nikolaidis, et al. 2008), in which the CDRL1 and CDRL2 were excluded. According to Das et al., positional numbering of cladistic markers was maintained.

### Intergenic and V-intron distance analysis

A comparison of *A. mexicanum* V-V intergenic length was made with *D. rerio* (GRCz11) and *H. sapiens* (GRCh38.p12). In these cases, coordinate data was obtained from ENSEMBL gff dump (Cunningham et al. 2022). As for *X. tropicalis* (Xenopus_tropicalis_v9.1, GCA_000004195.3).

### Recombination signal sequence (RSS) identification and analysis

Forty-two base pairs downstream of each presumably functional Variable segment were aligned with Clustal X and manually edited according to the vertebrate consensus CACAGTG and ACAAAAACC. A Positional weight matrix was calculated for the 7 and 9-mer, and the spacer length was calculated. A *bona fide* RSS was considered when the heptamer and nonamer weight score was ≥ half the corresponding maximal score and the spacer length was 23 ± 1 bp long (IGHV, IGHJ, IGLV, and IGSJ) or 12 ± 1 bp long (IGHD, IGLJ, and IGSV).

### Switch regions identification

We used the chr13q 35-50 Mbp interval to search in both strands for the occurrence of the AID hotspot motif 5’-AGCT-3’ as well as its iterations 5’-RGYW-3’ and 5’-WGCW (Xu et al. 2012) using DNA-Pattern at RSA tools (Medina-Rivera et al. 2015). Raw counts were estimated in 2500 bp non-overlapping windows. As count distribution is Gaussian, window counts were transformed to Z-scores to allow comparability between different motifs.

### Intron length analysis

V-intron length was calculated from the exon coordinates of the respective *locus* annotation file. In the case of human and zebrafish, coordinate data were retrieved using BioMart from ENSEMBL (http://www.ensembl.org/biomart/martview/). Due to the presence of abnormally long introns (outliers) in *A. mexicanum*, a robust ANOVA One-Way Trimmed Means Comparison with a trimming level of 5% (tr = 0.05), followed by the *post hoc* Lincon test was performed (Mair and Wilcox 2020). To calculate the enrichment of non-functional V genes according to intron length, a Fisher’s Exact test was implemented in R.

### Whole-genome, IGHV, and *p450* intergenic length analysis

Amex_v6 gene annotation file, Biomart, Kruskal-Wallis test with *post hoc* Dunn test correction for multiple comparisons implemented in R.

### Spleen RNA obtention for antibody heavy V region repertoire analyses

Three *A. mexicanum* adults specimens (two male and one female) were euthanized following institution and protocols with the authorization of the institutional board regulating and endangered species (SEMARNART: DGVS-PIMVS-CR-IN-1833/CDMX/17; N° SGPA/DGVS/04821). To generate libraries of *A. mexicanum* of variable regions of IgM, we synthesized cDNA using a modification of the method described (Matz et al. 1999). For cDNA synthesis, one microgram of RNA was purified by Trizol from the spleen. The IlluRACE oligonucleotide was used as template switching oligo (Additional file 2: Table S7) for the SuperScript II Reverse Transcriptase (Invitrogen). The amplicon libraries were amplified by PCR-RACE 5’ using oligonucleotides complementary to the CH1 exon in the heavy chain constant regions IgM of *A. mexicanum* in combination with the FpAmpIllu oligonucleotide (Additional file 2: Table S7). The CH1 oligonucleotide contains the reverse adapter of the Illumina MiSeq platform. The amplified libraries were gel purified, quantified by fluorometry, and subsequently re-amplified (Second stage PCR or index PCR) using the Nextera XT Index kit according to the 16S Metagenomic Sequencing Library instructions Preparation MiSeq Illumina protocol. Library sequencing was performed using an Illumina MiSeq platform in paired-end mode (2 × 250, 500 cycles).

## Acknowledgements

We would like to thank Dr. Sibayshi Das for sharing *X. tropicalis* V gene sequences; Drs. Sergei Nowoshilow and Jeramiah Smith for helpful discussions regarding scaffolding and chromosome mapping of the axolotl genome; Dr. Robert D. Miller for helpful discussions; Drs. Carolina González Torres and Francisco Javier Gaytán Cervantes for sequencing support (sequencing facility, Hospital de Especialidades Centro Médico Nacional Siglo XXI. IMSS); and Edgar Aguilar Vera and Everardo Gutierrez (Instituto Nacional de Salud Pública) for bioinformatics support. SSRH (CVU: 921827) and DLPO (CVU: 890789) are recipients of CONACyT scholarship.

## Contributions

JMB conceived and designed the study and contributed to data analysis, interpretation and manuscript writing. CLM conceived, designed the study and manuscript writing. EEGL contributed to data collection, analysis and interpretation. SSRH and DLPO contributed to data acquisition, analysis and drafting the manuscript. JMTS and HVT contributed to Rep-seq data acquisition and analysis. LZG and HMG provided and selected axolotl for the study, as well as training for animal handling, organ identification, and sample collection. RPP contributed to axolotl organ collection and samples processing. All authors reviewed and approved the final version.

## Notes

### Competing Interest Statement

The authors have declared no competing interest.

