## Additional file 1 for "Subfunctionalization and constrained size of the immunoglobulin *loci* in *Ambystoma mexicanum*"

### Supplementary figures

### Index

**Figure S1:** Schematic representation of IGH *locus* in *A. mexicanum* genome (v6) and the corresponding locus in *X. tropicalis.*
**Figure S2:** Representation of the major IGHV cluster in *A. mexicanum* genome (v6) in chr13q.
**Figure S3:** Density of AID hotspot motifs along the IGHC cluster.
**Figure S4:** Characterization of RGYW motif composition in the Sμ, Sχ and Sυ regions.
**Figure S5:** Analysis of the structure of the *A. mexicanum* IGHJ segments.
**Figure S6:** Structural analysis of IGHD segments within the 36.6 - 36.8 Mbp interval in chr13q.
**Figure S7:** Phylogenetic tree of human, mouse, and axolotl functional IGHV genes.
**Figure S8:** IGHV families.
**Figure S9:** Comparative analysis of the Constant Ig domain of light chains in humans and amphibian.
**Figure S10:** Structural analysis of *A. mexicanum* IGLJ segments.
**Figure S11:** Structural analysis of *A. mexicanum* IGSJ segments.
**Figure S12:** Cladistic analysis of light chain *loci.*
**Figure S13:** Mean distance phylogenetic tree of *A. mexicanum* Vλ protein sequences.
**Figure S14:** Schematic representation of immunoglobulin lambda *locus* in *A. mexicanum* genome.
**Figure S15:** Schematic representation of Immunoglobulin sigma *locus* in *A. mexicanum* genome.
**Figure S16:** Absence of the Ig Kappa *locus* in the *A. mexicanum* genome (v6).
**Figure S17:** IGHM gene length is evolutionarily constrained.
**Figure S18:** Functionality according to the IGH and IGL *loci* intron length.

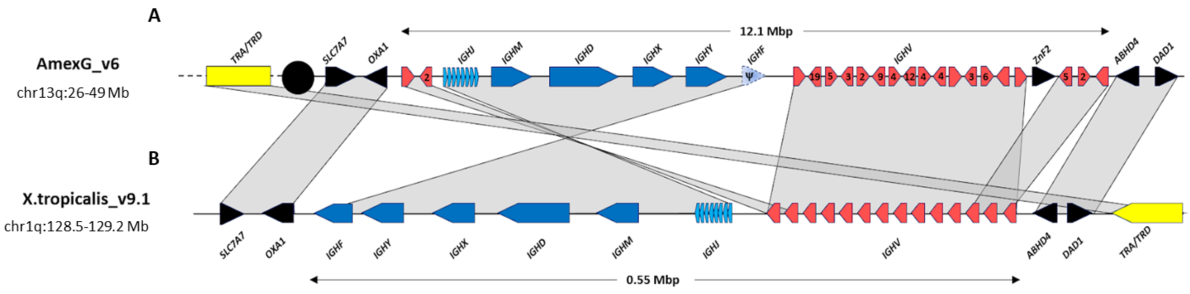


**Figure S1: Schematic representation of IGH *locus* in A) *A. mexicanum* genome (v6) and the corresponding *locus* in B) *X. tropicalis*.** The constant region cluster is shown in dark blue, J cluster in light blue, and V cluster in red, regardless of their functionality. Numbers depict the number of IGHV segments in a given orientation. Non-Ig genes in black. *X. tropicalis* displays the canonical architecture, in which the V, J and C clusters have the same orientation. Note that in *A. mexicanum* the JH-CH cluster and 3 IGHV segments appear to be inverted, which seems implausible due to the mechanism of V(D)J recombination. Moreover, IGHV segment orientation in axolotl are intercalated, which is atypical. Interestingly, in axolotl, the TRA/TRD *locus* (Yellow) is not linked to the IGH *locus* as in *X. tropicalis* and is coded across the centromere (black circle) in chr13p: 3.3-7.7 Mbp). Not on scale. The IGHD cluster is not shown for simplicity.

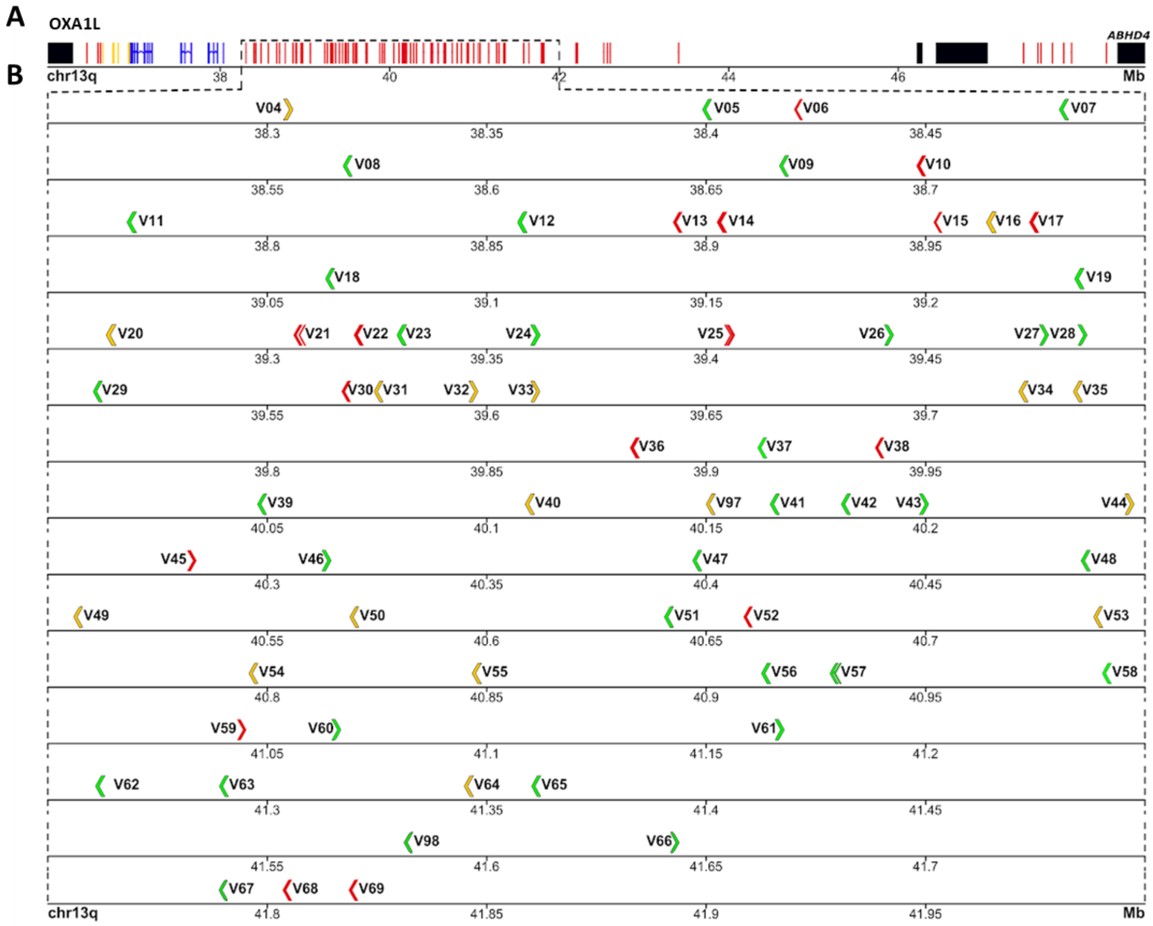


**Figure S2: Representation of the major IGHV cluster in *A. mexicanum* genome (v6) in chr13q**. **A)** IGH *locus* (chr13q: 35.96 - 48.91 Mbp). Genes in black are non-Ig genes, IGHC in blue, IGHV genes in red, and IGHJ and IGHD genes in yellow. **B)** Zoomed region of the main IGHV gene cluster (chr13q: 38.25- 42 Mbp). IGHV genes and their corresponding orientation are shown in arrows. Functional genes are in green, ORF's in yellow, and pseudogenes in red.

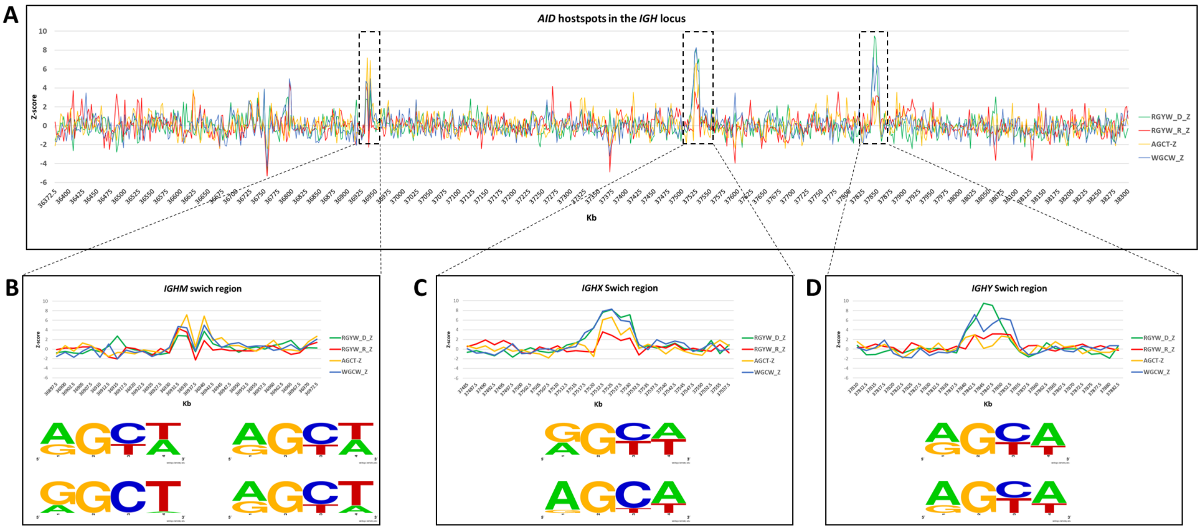


**Figure S3: Density of AID hotspot motifs along the IGHC cluster**. The frequency of each motif distribution per 2.5 Kb was used to calculate the corresponding Z-score. The RGYW in the direct strand is shown in green. RGYW in the opposite strand is shown in red. The palindromic AGCT and WGCW motifs are shown in yellow and blue, respectively. **A)** Panoramic of the IGHC cluster (chr13q: 36,732.5-38,300.0 Kb). **B)** Close-up to the Sμ region. Note that the central region is depleted of RGYW motifs giving an "M" shape. The logos at the bottom represent frequency per site for each of two Sμ peaks. Top row (direct strand), bottom row (reverse strand). **C)** Close-up to the Sχ region and its corresponding logos (direct, Top; reverse, bottom). **D)** Close-up of the Sυ region and its corresponding motif logos (direct, top; reverse, bottom)

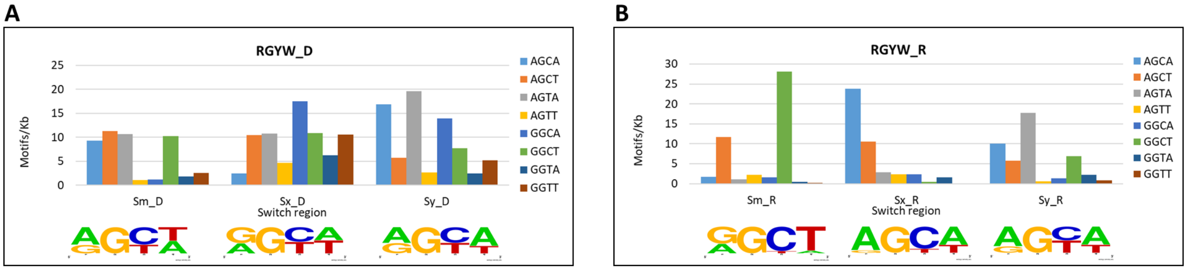


**Figure S4: Characterization of RGYW motif composition in the Sμ, Sχ and Sυ regions.** The frequency per Kb of the 8 tetramers corresponding to the RGYW motif is shown per switch region. For Sμ, the RGYW-depleted central region (2.5Kb) was excluded from the computation. **A)** The direct strand, with the corresponding frequency per site plot; and **B)** The reverse strand, with the corresponding frequency per site plot.

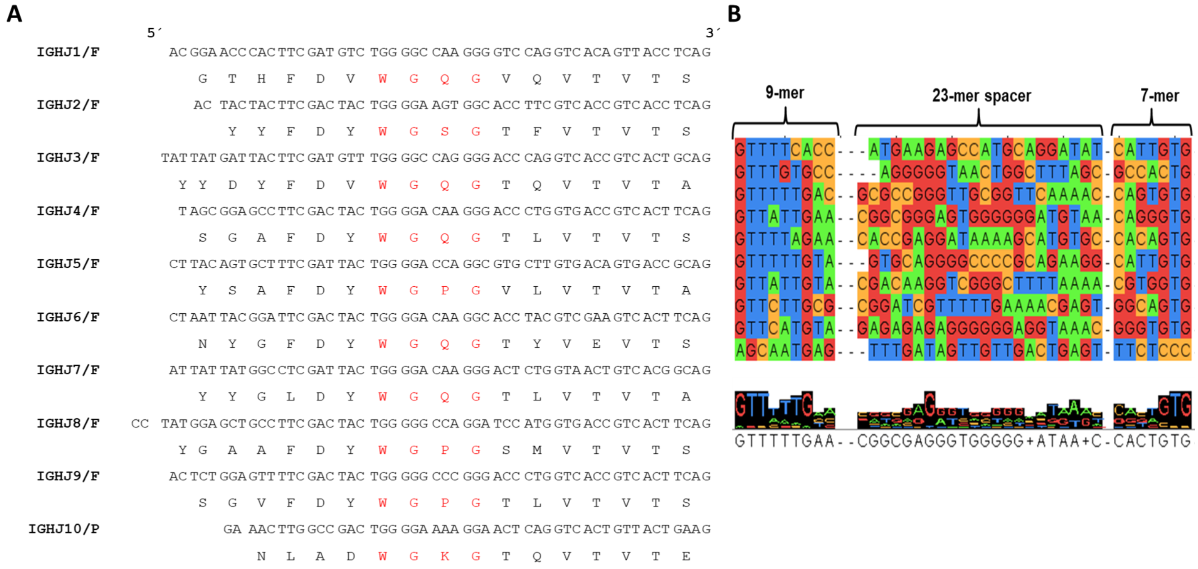


**Figure S5: Analysis of the structure of the *A. mexicanum* IGHJ segments.** The Axolotl IGHJ cluster comprises ten mapped genes, of which nine are functional, and one is a pseudogene (IGHJ10). This cluster is located on chromosome 13q, downstream of the IGHJ *locus*. **A)** Alignment of nucleotide and amino acid sequences of the ten identified JH segments, the conserved WGXG motif, characteristic of the IGH J-REGION, is in red. **B)** Alignment of nucleotide sequences of the RSS showing the conserved heptamer and nonamer separated by a 23 spacer. Consider that the sequences shown correspond to the direct chain 5'-3'.

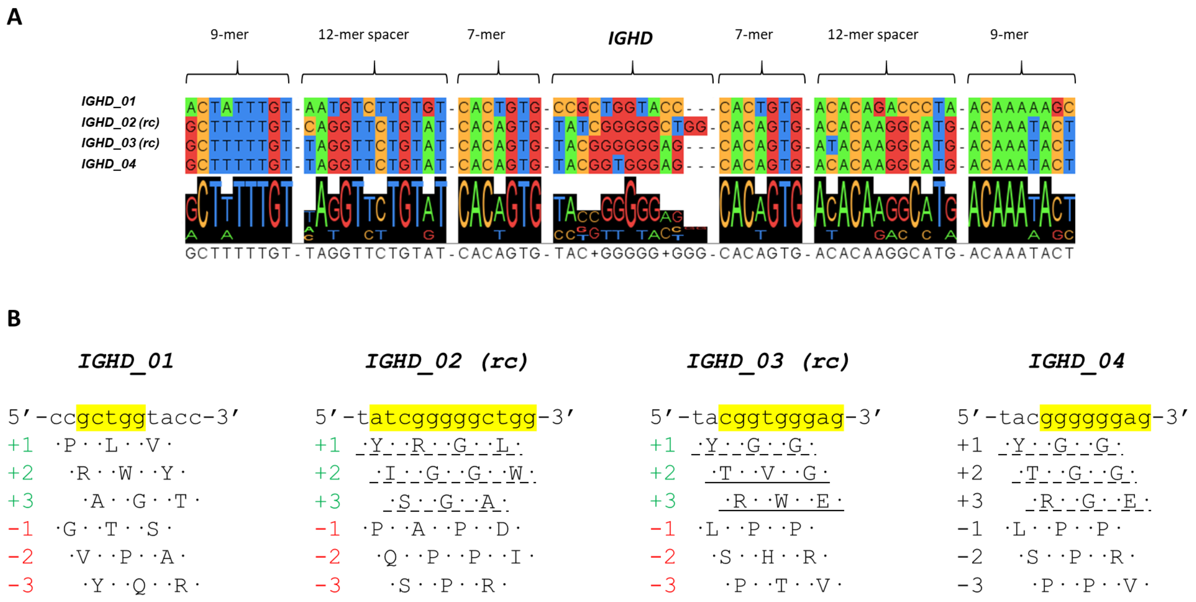


**Figure S6: Structural analysis of IGHD segments within the 36.6 - 36.8 Mbp interval in chr13q**. **A)** Multiple alignments of the four identified IGHD plus the RSS's in both flanks show the conserved 9 mer and 7 mer separated by a 12 mer spacer on both flanks. Note that in the genome assembly, IGHD_02 and IGHD_03 are in reverse orientation so that the reverse complement (rc) is shown. **B)** Sequence translation of IGHD segments. The nucleotide sequence is shown in lower case. Identical positions to DH-like core sequences described by Golub,1997 are highlighted in yellow (IGHD_01 = DH1, IGHD_02 = DH4, IGHD_03 = DH2, IGHD_04 = DH3). Translation in six reading frames is shown in uppercase letters. Translation frames that are found in our spleen Rep-seq dataset are solid underlined, whereas translation frames shared in Golub et al. and our spleen Rep-seq dataset are dotted underlined. IGHD_01 translations were not found in either dataset. Note that although all translation frames are productive, only forward translations were found in the peripheral repertoire in both datasets.

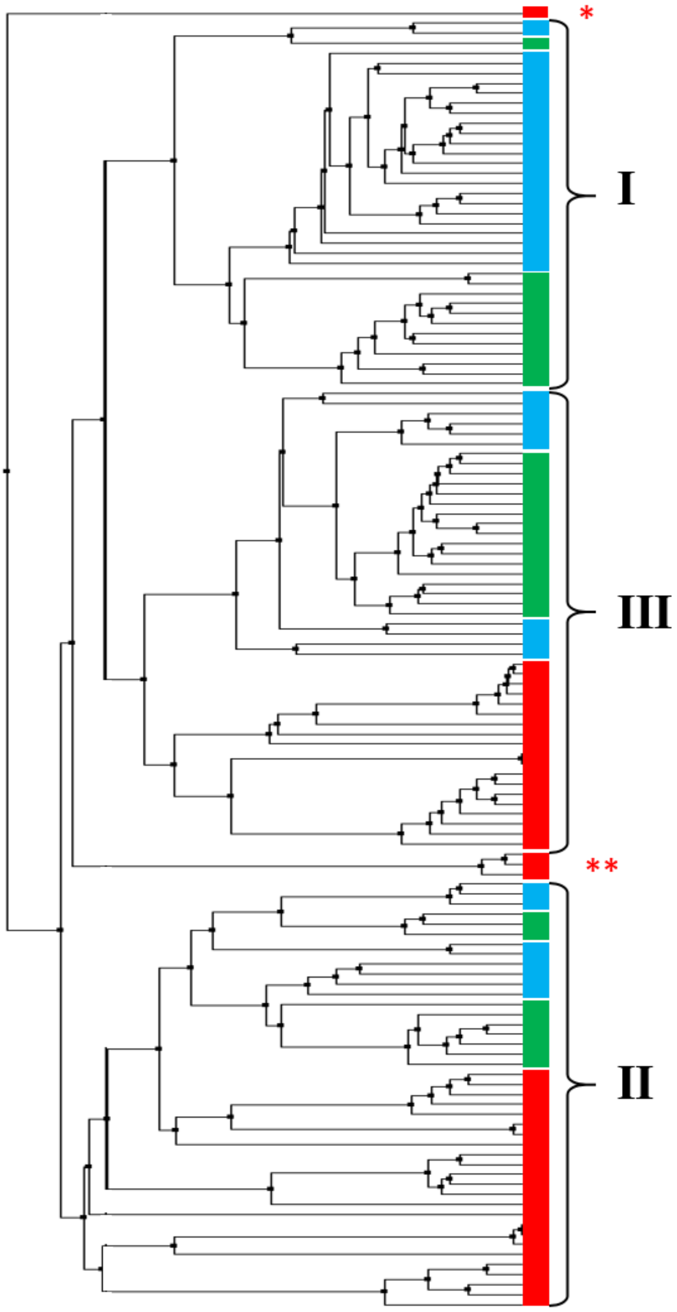


**Figure S7: Phylogenetic tree of human, mouse, and Axolotl functional IGHV genes.** Representative Human (Green bar) and mouse (light blue bar) sequences for the three tetrapod IGHV clans were used as references to classify *A. mexicanum* IGHV sequences (Red bar). No clan-I IGHV sequences were found in *A. mexicanum*. Note that IGHV_082 is atypical and outroots the tree (*). Nevertheless, IGHV_082 qualifies as a functional gene because it contains a Variable Ig domain ORF including Cys23 and Cys104, putatively functional RSS, and transcribed. Another atypical case (**) corresponds to IGHV_70, 071, and 077, which out root from the clan I/III stem, so we could not assign them to either clan.

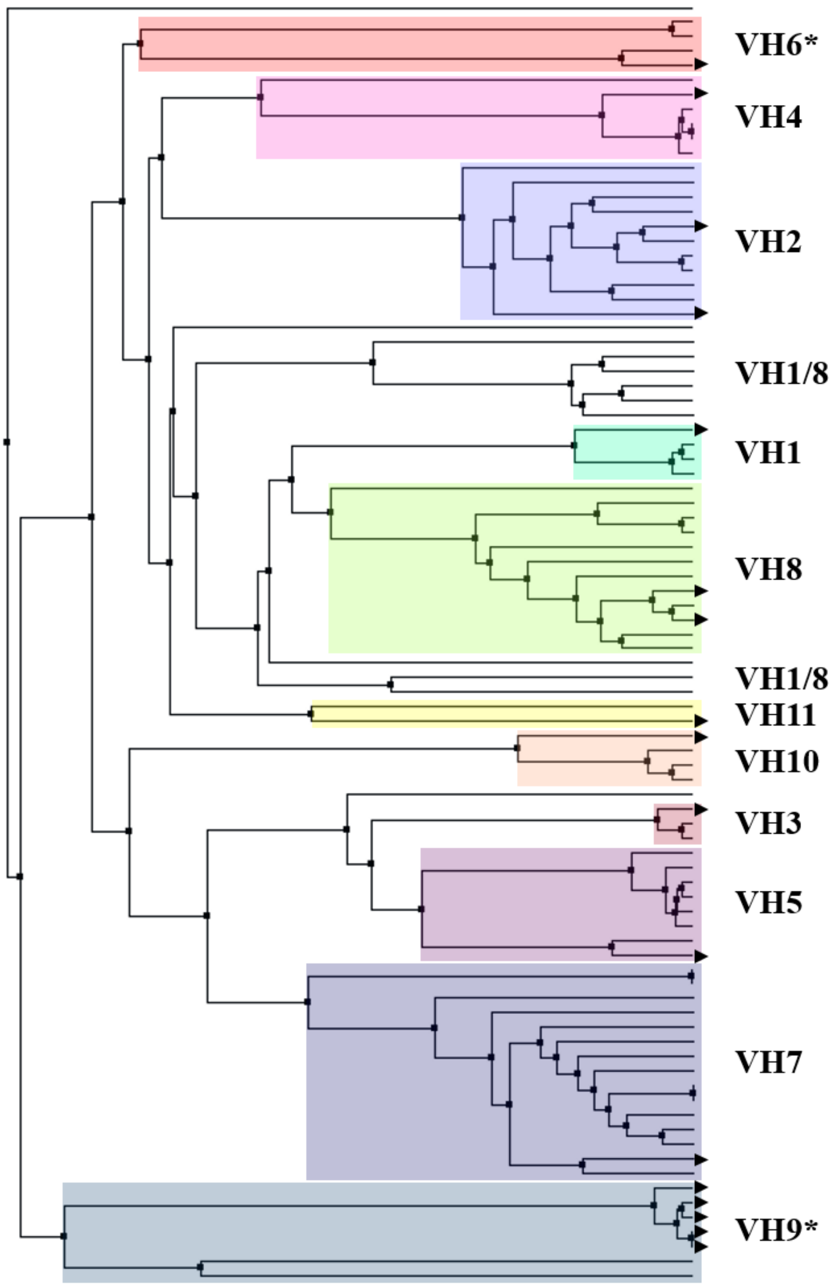


**Figure S8: IGHV families.** Classification of *A. mexicanum* germline functional and ORF IGHV genes according to the 11 families described by Golub et al., based on axolotl VH cDNA analysis (black triangles). Germline sequences and representative cDNA sequences of each family used as reference were aligned with MUSCLE, and the phylogenetic tree was based on mean protein distance. No functional IGHV genes belonged to VH family VH6 and VH9 (*). Germline sequences with white background correspond to sequences that could not be assigned to a particular family.

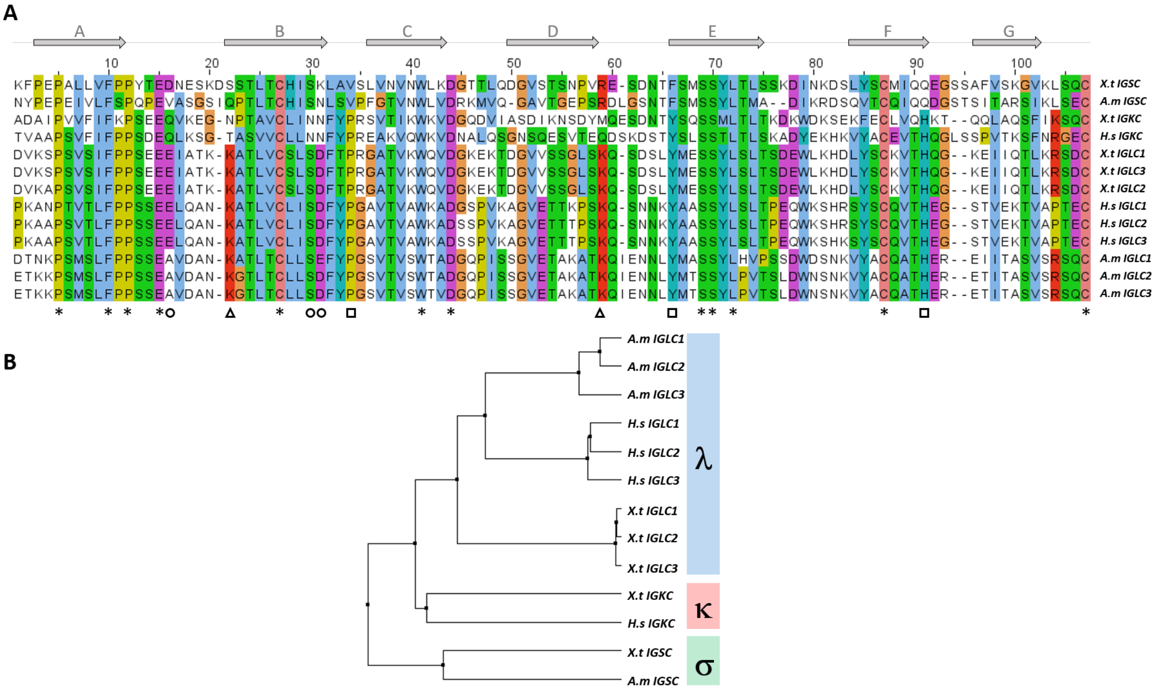


**Figure S9: Comparative analysis of the Constant Ig domain of light chains in humans and amphibians. A)** Multiple alignments of Cλ in *A. mexicanum*, *X. tropicalis* and human (only 3 out of 5 functional Cλ), Cσ in *A. mexicanum* and *X. tropicalis,* and Cκ in *X. tropicalis* and human. The seven canonical β-sheets (A-G) are shown on top of the alignment. Asterix shows twelve conserved residues in the three species, including Cys28, Trp42, Cys87, and Cys105. Some of these residues are involved in contact with H chains, whereas Cys28 and Cys87 mediate intradomain disulfide bonding, and Cys105 forms a disulfide bond with the H chain. Triangles show Lys23 and Lys60, which are unique to Cλ. Squares indicate Pro33, Tyr65, and His90, which are conserved in Cλ and Cκ, but are replaced by Val, Phe, and Gln, respectively in Cs. Circles show Gln17, Asp30, and Asp31 that are unique to Cκ. Alignment color is based on Clustal X coloring scheme. **B)** Average distance phylogenetic tree of Ig Light chain C domain based on the multiple alignments shown in A. The C-type domain clusters together regardless of the species.

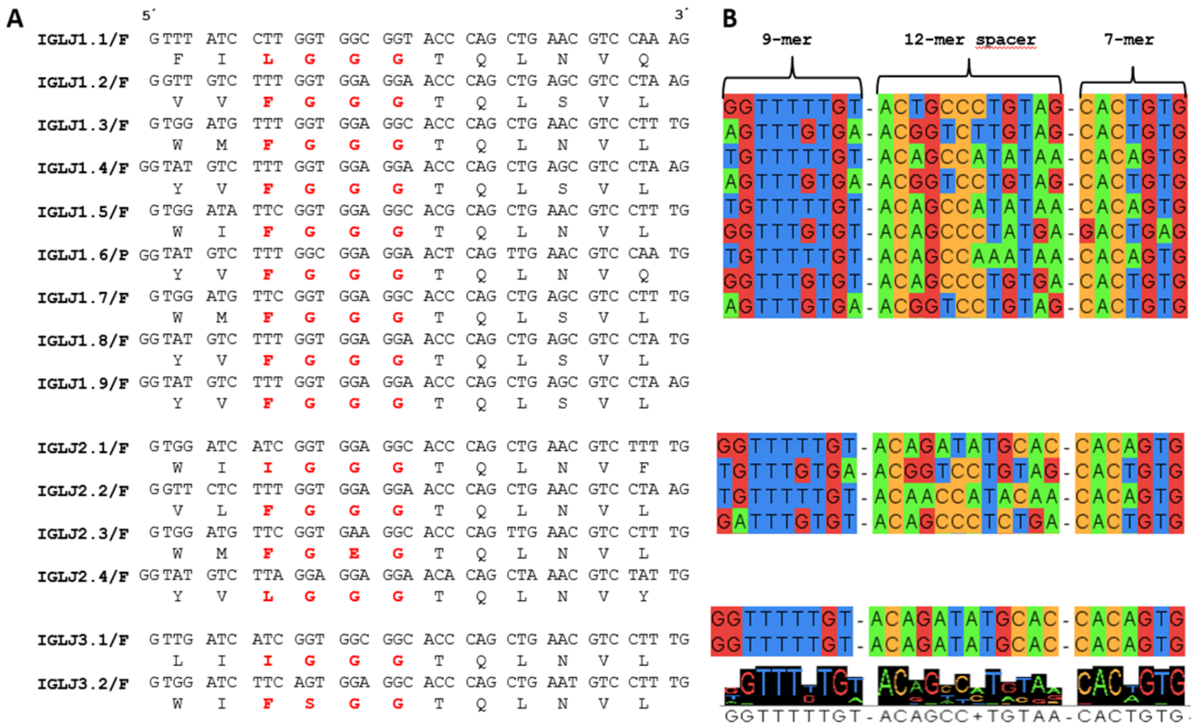


**Figure S10: Structural analysis of the *Ambystoma mexicanum* IGLJ segments.** The Axolotl IGLJ group comprises three clusters. The first cluster includes nine mapped genes, of which eight are functional, and one is a pseudogene (IGLJ6). The second cluster comprises four mapped functional genes. These groups are located on chromosome 10q, flanked by the *DGCR2* gene on the centromeric flank and by the *CABIN1*. **A)** Alignment nucleotide and amino acid sequences of the ten identified IGLJ segments, the conserved FGXG motif, characteristic of the IGL J-REGION, is in red. **B)** Alignment of nucleotide sequences of the RSS showing the conserved heptamer and nonamer separated by a 12 pb spacer. Considering that the sequences shown correspond to the direct chain 5'-3'.

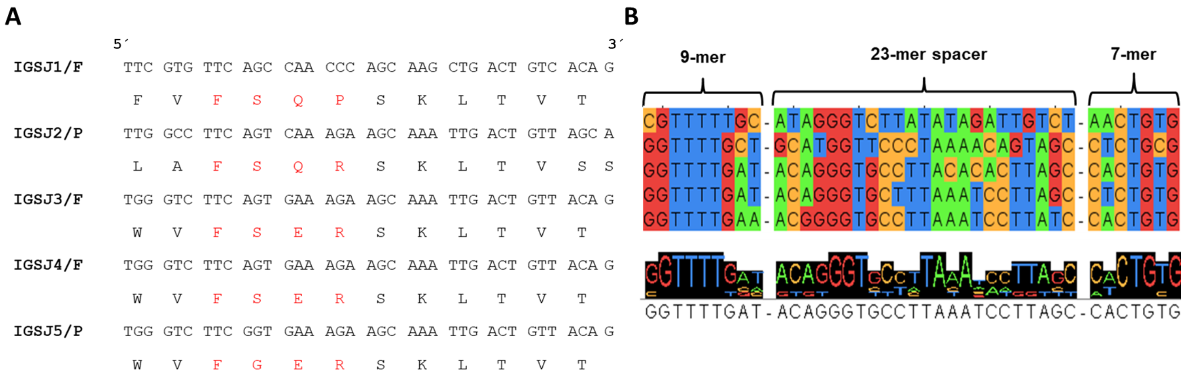


**Figure S11: Structural analysis of the *Ambystoma mexicanum* IGSJ segments.** The Axolotl IGSJ group comprises three functional segments and two pseudogenes (IGSJ2-IGSJ5). The IGSJ cluster is in the Chr1p. **A)** Aligning nucleotide and amino acid sequences of the five IGSJ segments, the conserved FSXR motif characteristic of the IGS J-REGION is in red. **B)** Alignment of nucleotide sequences of RSS showing the conserved heptamer and nonamer separated by a 23 pb spacer. Consider that the sequences shown correspond to the direct chain 5’-3'.

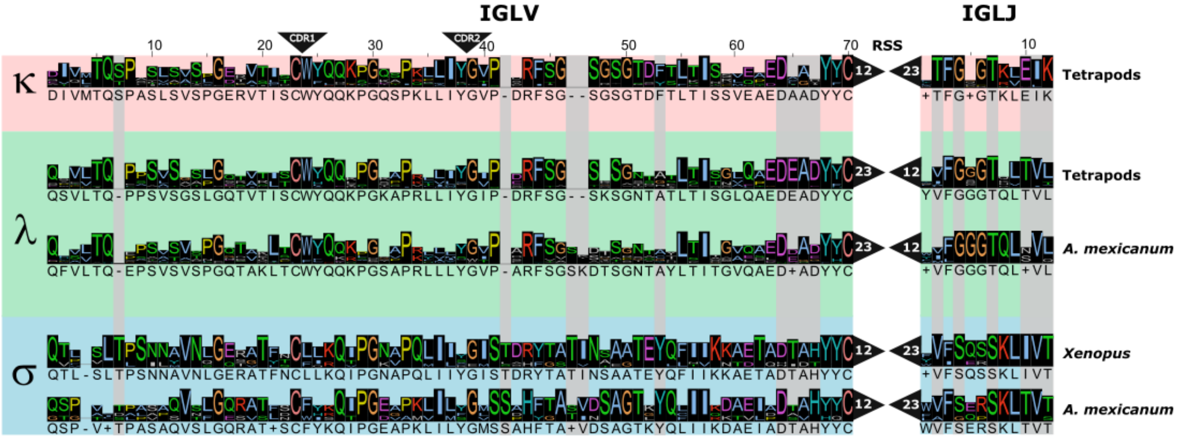


**Figure S12: Cladistic analysis of light chain *loci*.** Light chain protein sequences of tetrapods (V and J) were provided by S. Das, aligned with MUSCLE. Functional sequences of *A. mexicanum* were also included. The sequences corresponding to the CDRλ1 and 2 were removed. According to Das et al., 2008, Cladistic markers are shown in gray. Note that the position of markers is according to Das *et al*. (Without CDR1 and 2).

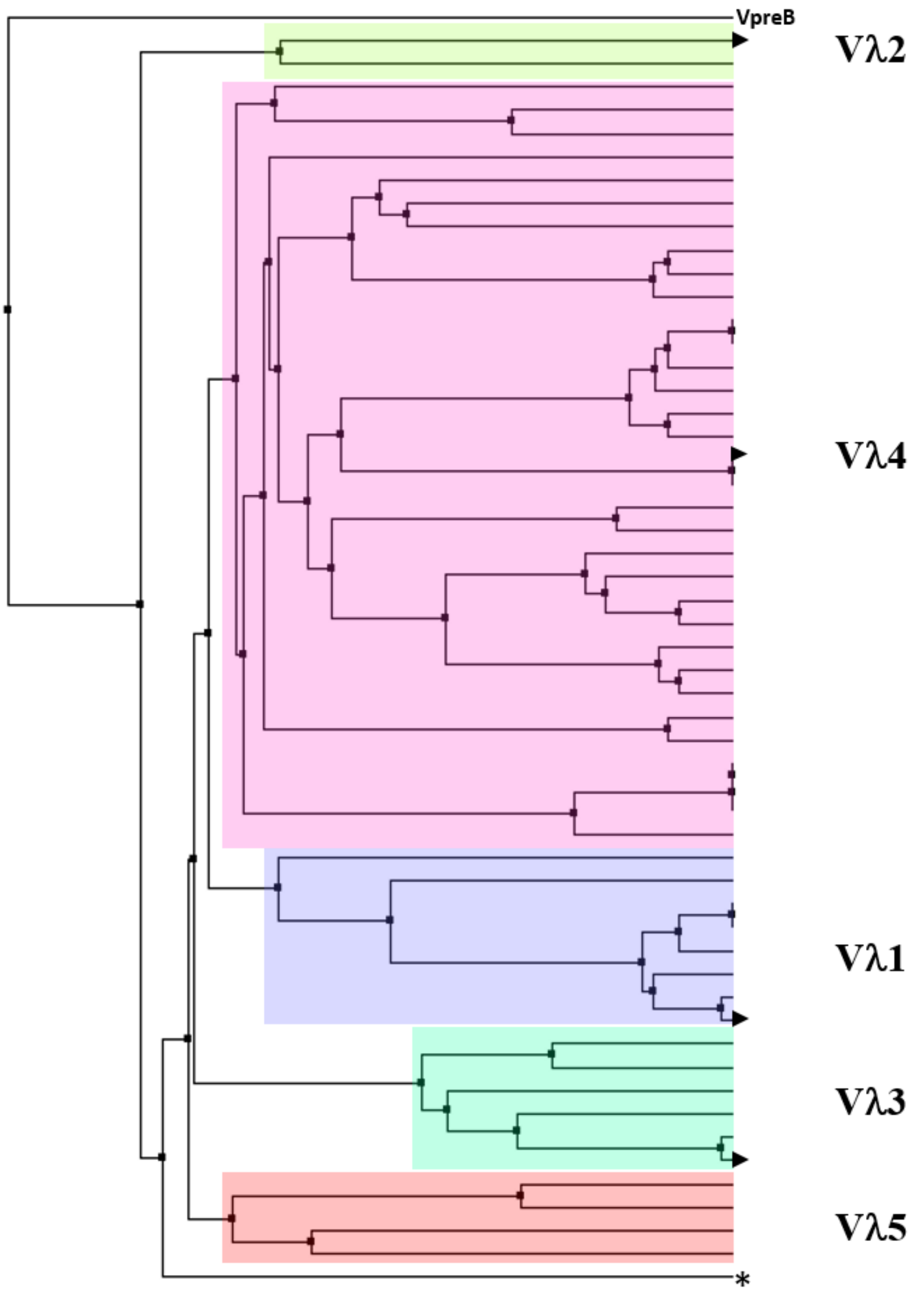


**Figure S13: Mean distance phylogenetic tree of *A. mexicanum* Vλ protein sequences.** We used V1, V2, V3, and V4 sequences described by Golub as references (black triangles) to classify Vl into previously described families (Accessions: AF317321.1, AF317322.1, AF317323.1, AF317324.1). VpreB protein sequence was included to root the tree. We found functional Vλ genes in the four families, but Vl4 was the most abundant (n = 29). Vλ2 had only one Vλ (IGLV_059). IGLV_040, IGLV_069, and IGLV_037 formed a previously unidentified family (Vl5). IGLV_039 did not cluster with any family (*).

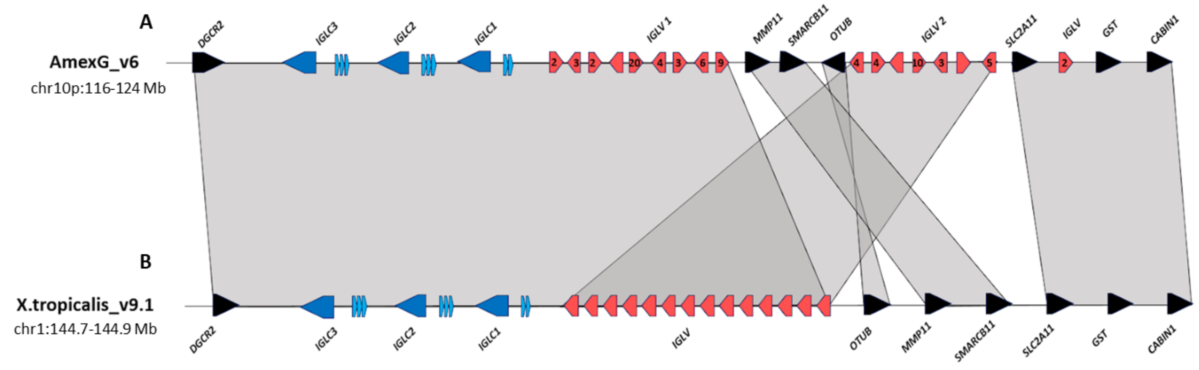


**Figure S14: Schematic representation of immunoglobulin lambda locus in A) A. mexicanum genome.** Located in chr10p:116-124 Mbp (v6), and **B)** *X. tropicalis* in chr1:144.7-144.9 Mbp. The constant region clusters are shown in dark blue, J clusters in light blue, and V clusters in red, regardless of their functionality. Numbers depict the number of IGLV segments in a given orientation. Non-Ig genes in black. *X. tropicalis* displays the canonical architecture, in which the V, J, and C clusters are in the same orientation. Note that in *A. mexicanum,* IGLV genes are in at least two different clusters (IGLV1 and IGLV2), separated by the *MMP11*, *SMARCB11,* and *OTUB* genes, which in *Xenopus* are located in the distal flank. Also, note that the *OTUB* gene in axolotl is inverted. Moreover, as in the IGHV *locus,* IGLV segment orientation in axolotl is intercalated, which is atypical. VJ recombination involving the more distal IGLV segments would delete the *OTUB*, *SMARCB11*, *MMP11,* and the *SLC2A11* gene cluster. Not on a scale.

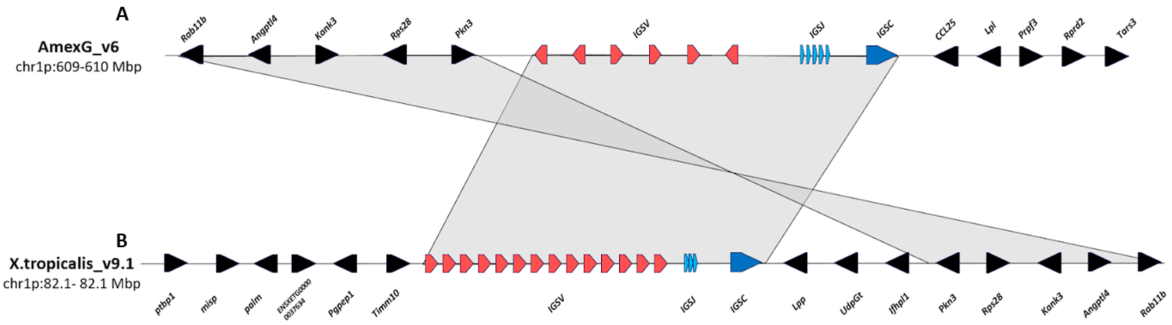


**Figure S15: Schematic representation of Immunoglobulin sigma *locus* in A) *A. mexicanum* genome (v6).** Located in chr1p:609-610 Mbp (v6), and **B)** *X. tropicalis* in chr1: 82.1 - 82.22 Mbp. The constant region clusters are shown in dark blue, J clusters in light blue, and V clusters in red, regardless of their functionality. Non-Ig genes in black. *X. tropicalis* displays the canonical architecture, in which the V, J and C clusters are in the same orientation. Note that in *A. mexicanum, IGSV* gene orientation in axolotl is intercalated, which is atypical. Non-Ig genes in the downstream flank are different in *A. mexicanum* and at least the *Prpf3*, *Rprd2* and *Tars3* genes are located in Chr8 106.9 – 107.1 Mbp in *X. tropicalis*. Also, non Ig genes in the upstream flank correspond to the downstream flank in *X. tropicalis*. It is possible that this chromosomal configuration is correct, however it is also possible that the *IGS locus* may be inverted so that the downstream flank is syntenic with *X. tropicalis*. Not on scale.

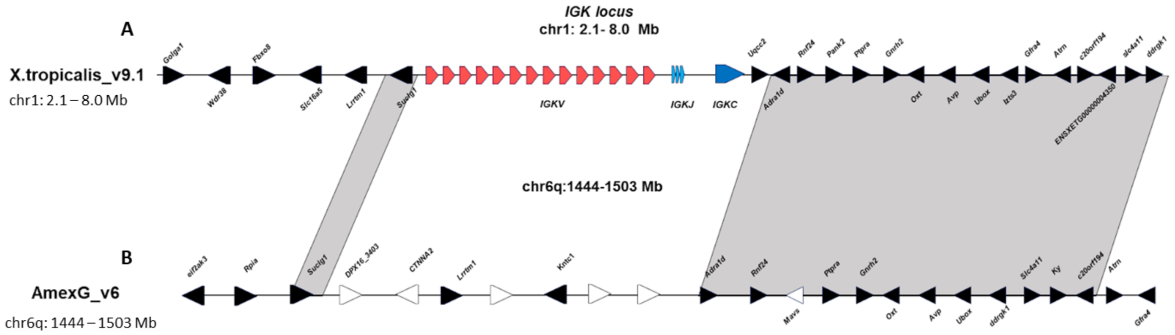


**Figure S16: Absence of the Ig kappa *locus* in the *A. mexicanum* genome (V6).** Schematic representation of the **A)** *X. tropicalis* κ *locus* in chr1: 2.1 – 8.0 Mbp. The constant region gene is shown in dark blue, J cluster in light blue, and V cluster in red, regardless of their functionality. Non-Ig genes in black. *X. tropicalis* displays the canonical architecture, in which the V, J, and C clusters are in the same orientation. Note that in **B)** *A. mexicanum,* there is a complete absence of the kappa *locus,* and the downstream flank is located in chr6q: 1444 – 1503 Mbp. Two 1:1 orthologs in the upstream flank, *LRRTM1,* and *SUCLG1,* also map to the same region. Non-orthologous genes are shown as white-filled genes. Not on a scale.

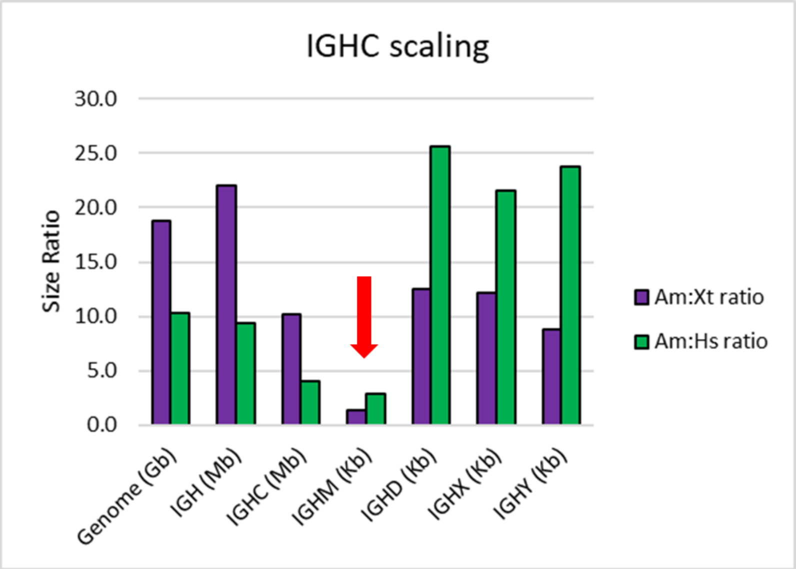


**Figure S17: IGHM gene length is evolutionarily constrained.** *A.mexicanum* to *X. tropicalis* (purple bars) or *H. sapiens* (green bars) ratios for the whole genome, IGH *locus* (from the first IGHV gene to the last IGHC gene), IGHC *locus* (from the first to last IGHC gene), and individual IGHC genes (From first to the last exon). To calculate IGHC ratios in humans, the human orthologs for IGHX (IGHG) and IGHY (IGHA) were used.

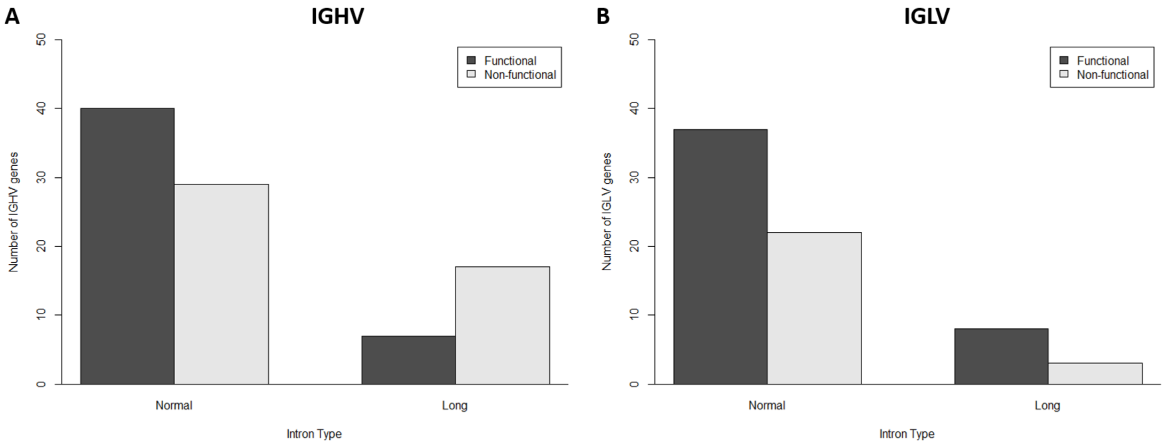


**Figure S18: Functionality according to the A) IGH and B) IGL *loci* intron length.** A 2 x 2 contingency table was built with the number of functional and non-functional (pseudogenes plus ORF's) according to V-intron length (long V-intron > 150 bp) for IGHV (left) and IGLV (right). A Fisher's exact test revealed that the odds of an IGHV gene with a long V-intron being non-functional are 3.3 higher than its short V-intron counterpart (P = 0.018, CI95: 1.1, 10.7). In contrast, for the IGLV *locus,* there were no differences in the observed and expected frequencies of short and long V-introns according to functionality (P = 0.73; OR = 0.63, CI95: 0.09, 3.0).
